# Mechanisms underlying divergent responses of genetically distinct macrophages to IL-4

**DOI:** 10.1101/2020.11.02.365742

**Authors:** Marten A. Hoeksema, Zeyang Shen, Inge R. Holtman, An Zheng, Nathan Spann, Isidoro Cobo, Melissa Gymrek, Christopher K. Glass

## Abstract

Mechanisms by which non-coding genetic variation influences gene expression remain only partially understood but are considered to be major determinants of phenotypic diversity and disease risk. Here, we evaluated effects of >50 million SNPs and InDels provided by five inbred strains of mice on the responses of macrophages to interleukin 4 (IL-4), a cytokine that plays pleiotropic roles in immunity and tissue homeostasis. Remarkably, of >600 genes induced >2-fold by IL-4 across the five strains, only 26 genes reached this threshold in all strains. By applying deep learning and motif mutation analyses to epigenetic data for macrophages from each strain, we identified the dominant combinations of lineage determining and signal-dependent transcription factors driving late enhancer activation. These studies further revealed mechanisms by which non-coding genetic variation influences absolute levels of enhancer activity and their dynamic responses to IL-4, thereby contributing to strain-differential patterns of gene expression and phenotypic diversity.

## Introduction

Non-coding genetic variation is a major driver of phenotypic diversity as well as the risk of a broad spectrum of diseases. For example, of the common single nucleotide polymorphisms (SNPs) and short insertions/deletions (InDels) identified by genome-wide association studies (GWAS) to be linked to specific traits or diseases, ~90% are typically found to reside in non-coding regions of the genome (Farh et al., 2015). The recent application of genome-wide approaches to define the regulatory landscapes of many different cell types and tissues allows intersection of such variants with cell-specific regulatory elements and strongly supports the concept that alteration of transcription factor binding sites at these locations is an important mechanism by which they influence gene expression (Kilpinen et al., 2013; van der Veeken et al., 2019; Vierstra et al., 2020). Despite these major advances, it remains difficult to predict the consequences of most forms of non-coding genetic variation. Major challenges that remain include defining the causal variant within a block of variants that are in high linkage disequilibrium, identifying the gene that is regulated by the causal variant, and understanding the cell type and cell state specific regulatory landscape in which a variant might have a functional consequence (Consortium et al., 2020). For example, a variant that affects the binding of a signal-dependent transcription factor (SDTF) may only be of functional importance in a cell that is responding to a signal that activates that factor (Soccio et al., 2015). Also, sequence variants can have a range of effects on transcription factor binding motifs, from abolishing binding or to changing it to a high affinity motif by affecting critical nucleotides, or by increasing/decreasing binding to an intermediate affinity motif by changing nucleotides that quantitatively affect binding (Behera et al., 2018; Deplancke et al., 2016; Grossman et al., 2017).

Studies of the impact of natural genetic variation on signal-dependent gene expression have demonstrated large differences in absolute levels of gene expression under basal and stimulated conditions, which result in corresponding differences in the dynamic range of the response (Bakker et al., 2018; Fairfax et al., 2014; Gate et al., 2018). The molecular mechanisms by which genetic variation results in these qualitatively and quantitatively different signal-dependent responses remain poorly understood but are likely to be of broad relevance to understanding how non-coding variation influences responses to signals that regulate development, homeostasis and disease-associated patterns of gene expression.

To investigate the influence of genetic variation on signal-dependent gene expression, we performed transcriptomic and epigenetic studies of the responses of macrophages derived from five different inbred mouse strains to the anti-inflammatory cytokine IL-4 (Figure 1A). The selected strains include both similar as well as highly divergent strain pairs, allowing modeling of the degree of variation between two unrelated individuals (~4 million variants) and that observed across large human populations (>50 million variants). Using this approach, we previously showed that strain-specific variants that disrupt the recognition motif for one macrophage lineage determining transcription factor (LDTF, e.g. PU.1), besides reducing binding of the LDTF itself, also result in decreased binding of other collaborative factors and SDTFs (Heinz et al., 2013; Link et al., 2018a). Collectively, these findings supported a model in which relatively simple combinations of LDTFs collaborate with an ensemble of additional transcription factors to select cell-specific enhancers that provide sites of action of broadly expressed SDTFs (Heinz et al., 2010).

**Figure 1.**
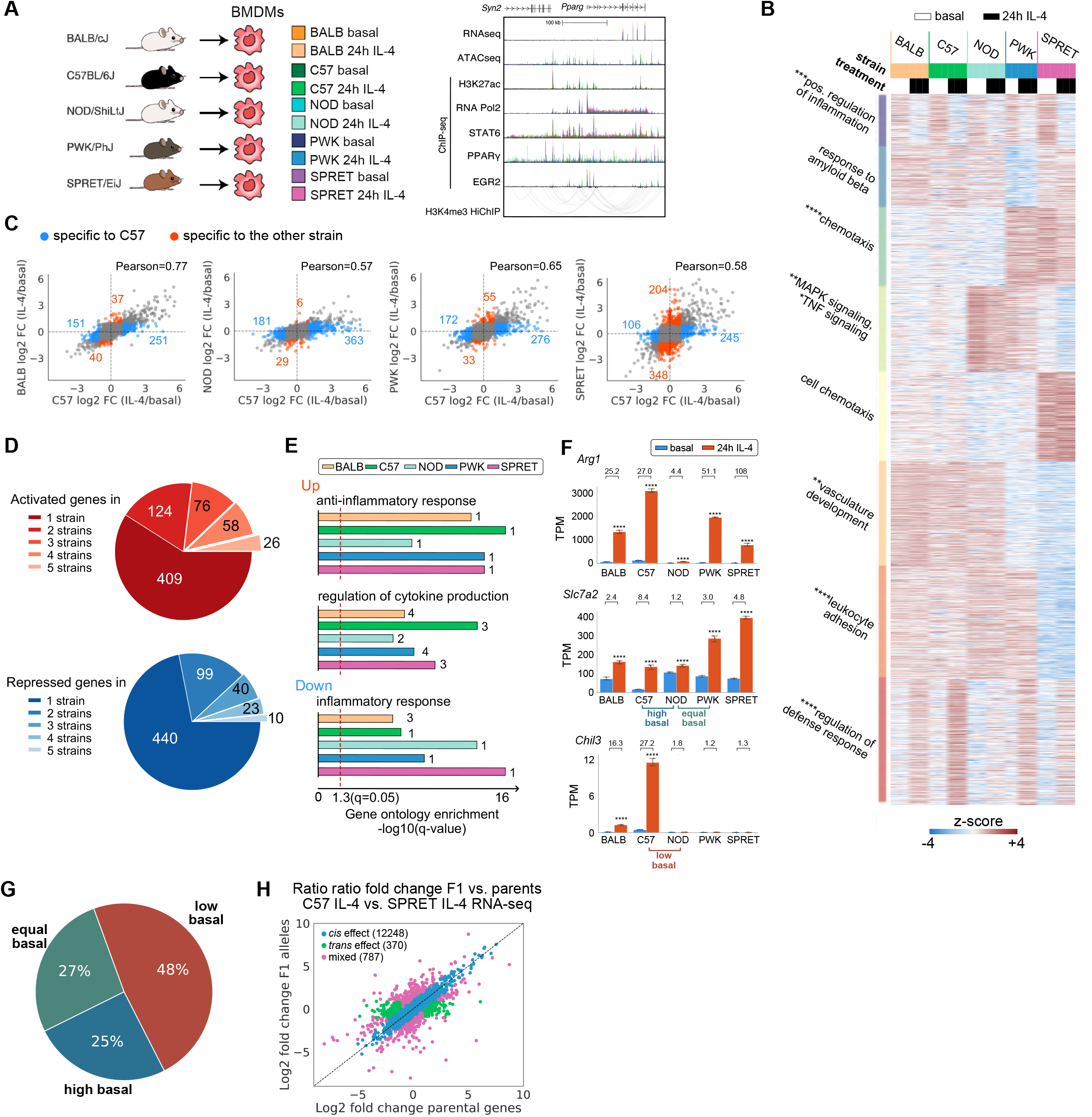
Response to IL-4 is highly divergent in BMDMs from different mouse strains. A. Overview of experimental design and main data sets. B. WGCNA clustering focused on strain-differentially regulated genes in IL-4 treated BMDMs. The top hit Metascape pathways are annotated for each module. *q<0.05, **q<0.01,***q<0.001, ****q<0.0001. C. Ratio-ratio plots demonstrating the mRNA response to IL-4 in pairwise comparisons. D. Overlap of genes significantly induced or repressed (q<0.05, >2-fold) after IL-4 treatment in BMDMs from all strains. E. Gene regulatory pathways up- and down-regulated after 24h IL-4 stimulation in BMDMs from all strains. Numbers indicate the rank order in pathway analysis. F. *Arg1, Slc7a2* and *Chil3* as example genes differentially up-regulated by IL-4 in strains. TPM, transcripts per kilobase million. ****q<0.0001, compared to basal. Numbers indicate fold change by IL-4. G. Effect of differences in basal gene expression on strain-differential IL-4 inductions. H. Average log2 gene expression fold change between alleles in hybrid (C57xSPRET F1) and parental strain under 24h IL-4 conditions.

IL-4 has many biological roles, including regulation of innate and adaptive immunity (Gieseck et al., 2018). In macrophages, IL-4 drives an ‘alternatively activated’ program of gene expression associated with inhibition of inflammatory responses and promotion of wound repair (Gordon and Martinez, 2010). The immediate transcriptional response to IL-4 is mediated by activation of STAT6 (Goenka and Kaplan, 2011; Ostuni et al., 2013), which rapidly induces the expression of direct target genes that include effector proteins such as Arginase 1 (*Arg1*) and transcription factors like PPARg (Daniel et al., 2018; Huang et al., 1999) and EGR2 (Daniel et al., 2020). However, the extent to which natural genetic variation influences the program of alternative macrophage activation has not been systematically evaluated. Here, we demonstrate highly differential IL-4 induced gene expression and enhancer activation in bone marrow-derived macrophages (BMDMs) across the five mouse strains, thereby establishing a robust model system for quantitative analysis of the effects of natural genetic variation on signal-dependent gene expression. Through the application of deep learning methods and motif mutation analysis of strain-differential IL-4 activated enhancers, we provide functional evidence for a dominant set of LDTFs and SDTFs required for late IL-4 enhancer activation, which include STAT6, PPARγ and EGR2, and validate these findings in *Egr2*-knockout BMDMs. Importantly, assessment of the quantitative effects of natural genetic variants on recognition motifs for LDTFs and SDTFs suggests general principles by which such variation affects enhancer activity patterns and dynamic signal responses.

## Results

### The response to IL-4 is highly variable in BMDMs from genetically diverse mice

To investigate how natural genetic variation affects the macrophage response to IL-4, we began by performing RNA-seq in BMDMs derived from female BALB/cJ (BALB), C57BL/6J (C57), NOD/ShiLtJ (NOD), PWK/PhJ (PWK) and SPRET/EiJ (SPRET) mice under basal conditions and following stimulation with IL-4. Time course experiments in C57 BMDMs indicated a progressive increase in the number of differentially expressed genes from 1 to 24 hours (Figure S1A-B, Table S1). We therefore focused our analysis on the response to IL-4 in BMDMs from the five strains at this timepoint. Weighted Co-expression Network Analysis (WGCNA) identified numerous modules of highly correlated mRNAs, the majority of which were driven by strain differences (Figure 1B). Genes that were positively regulated by IL-4 across strains (red module, bottom) were enriched for functional annotations related to negative regulation of defense responses. Conversely, the purple (top) module captured genes that were negatively regulated by IL-4 and were enriched for pathways associated with positive regulation of inflammation (Figure 1B).

Remarkably, of the 693 genes induced >2-fold in at least one strain, only 26 (3.75%) were induced at this threshold in all five strains (Figure 1D, S1C-D, Table S2). Conversely, more than half of the IL-4-responsive genes identified were induced >2-fold in only a single strain. NOD BMDMs were notable for a generally attenuated response to IL-4 (Figure 1B, red module, 1C, second panel). A similar pattern was observed for down-regulated genes (Figure 1D). Despite these differences at the level of individual genes, similar pathways/gene programs were enriched in all strains for both induced and repressed genes (Figure 1E). Substantial differences in IL-4 target gene expression across strains are illustrated by *Arg1, Slc7a2* and *Chil3* (Figure 1F). BMDMs from all strains exhibit a significant induction of *Arg1* expression, but the absolute basal levels and induction folds vary by more than an order of magnitude. *Slc7a2* exhibits similar levels of expression in C57 and NOD BMDMs after IL-4 treatment, but its differences at the basal level result in an 8-fold and 1.2-fold change, respectively. We refer to the pattern of reduced responsiveness to IL-4 in this comparison of C57 and NOD as being associated with ‘high basal’ activity in the less responsive strain. Conversely, NOD and PWK BMDMs exhibit similar levels of basal *Slc7a2* expression, but IL-4 only increased *Slc7a2* expression more than 2-fold in PWK. We refer to this pattern of reduced responsiveness to IL-4 in NOD compared to PWK as being associated with ‘equal basal’ activity. A third category is exemplified by *Chil3,* which is induced in C57 but not in NOD BMDMs. In this case, lack of responsiveness is associated with low expression of *Chil3* under basal conditions. We refer to this pattern as ‘low basal’ in the less responsive strain. Quantitative analysis of pair-wise comparisons indicate that 48% of the genes with decreased IL-4 induced gene expression were due to low basal expression, 27% had no differences prior to IL-4 stimulation (equal basal), and 25% were the result of a high basal expression level in the less responsive strain (Figure 1G).

To investigate local versus distant effects of genetic variation on the differential responses to IL-4, we crossed C57 mice with the most genetically distinct SPRET mice to generate F1 offspring containing each parental chromosome. 91.4% of parental-specific RNA-seq reads in the F1 strain are within 2-fold of their values in C57 and SPRET and considered to be due to local (*cis*) effects of genetic variation (Figure 1H, S1E), while only 2.8% was divergent in the F1 BMDMs, indicating *trans* regulation. Collectively, these studies uncovered striking variation in the cell autonomous responses of BMDMs to IL-4 across these five strains, providing a powerful experimental system for investigating mechanisms by which natural genetic variation impacts signal-dependent gene expression.

### Strain-differential IL-4 induced gene expression is associated with differential IL-4 enhancer activation

To investigate the impact of *cis* variation on putative transcriptional regulatory elements, we defined high confidence IL-4 activated enhancers as intronic or intergenic open chromatin regions (based on ATAC-seq) with at least 2.5-fold increase in H3K27ac (Creyghton et al., 2010) and RNA Pol2 (Bonn et al., 2012) after IL-4 treatment (Figure S2A-D). In 24-hour IL-4 stimulated C57 BMDMs, 1106 regions exhibited a >2.5-fold increase in H3K27ac, whereas 332 regions exhibited a >2.5-fold decrease, corresponding to putative IL-4-activated and IL-4-repressed enhancers, respectively (Figure 2A). Comparison of C57 enhancers to those of other strains under IL-4 treatment conditions revealed marked differences that scaled with the degree of genetic variation (Figure 2B, S2E-F). We further subdivided these regions into ‘conventional enhancers’ (blue, Figure 2C) and ‘super enhancers’ (orange, Figure 2C), based on the density distribution of normalized H3K27ac tag counts (Whyte et al., 2013). Super enhancers represent regions of the genome that are highly enriched for cell-specific combinations of transcription factors and coregulators and control the expression of genes required for cellular identity and critical functions. In comparison to conventional enhancers, super enhancers exhibited significantly less variation in H3K27ac in response to IL-4 (Figure 2D, S2G), suggesting relative resilience to genetic variation. For example, IL-4 induction of the *Ak2* super enhancer (Figure 2E) is highly conserved between the five strains. In contrast, a typical example of strain specificity is provided by the conventional enhancers associated with the *Msx3* gene. These enhancers are IL-4 inducible only in BALB, C57 and NOD and absent in PWK and SPRET BMDMs (Figure 2F).

**Figure 2.**
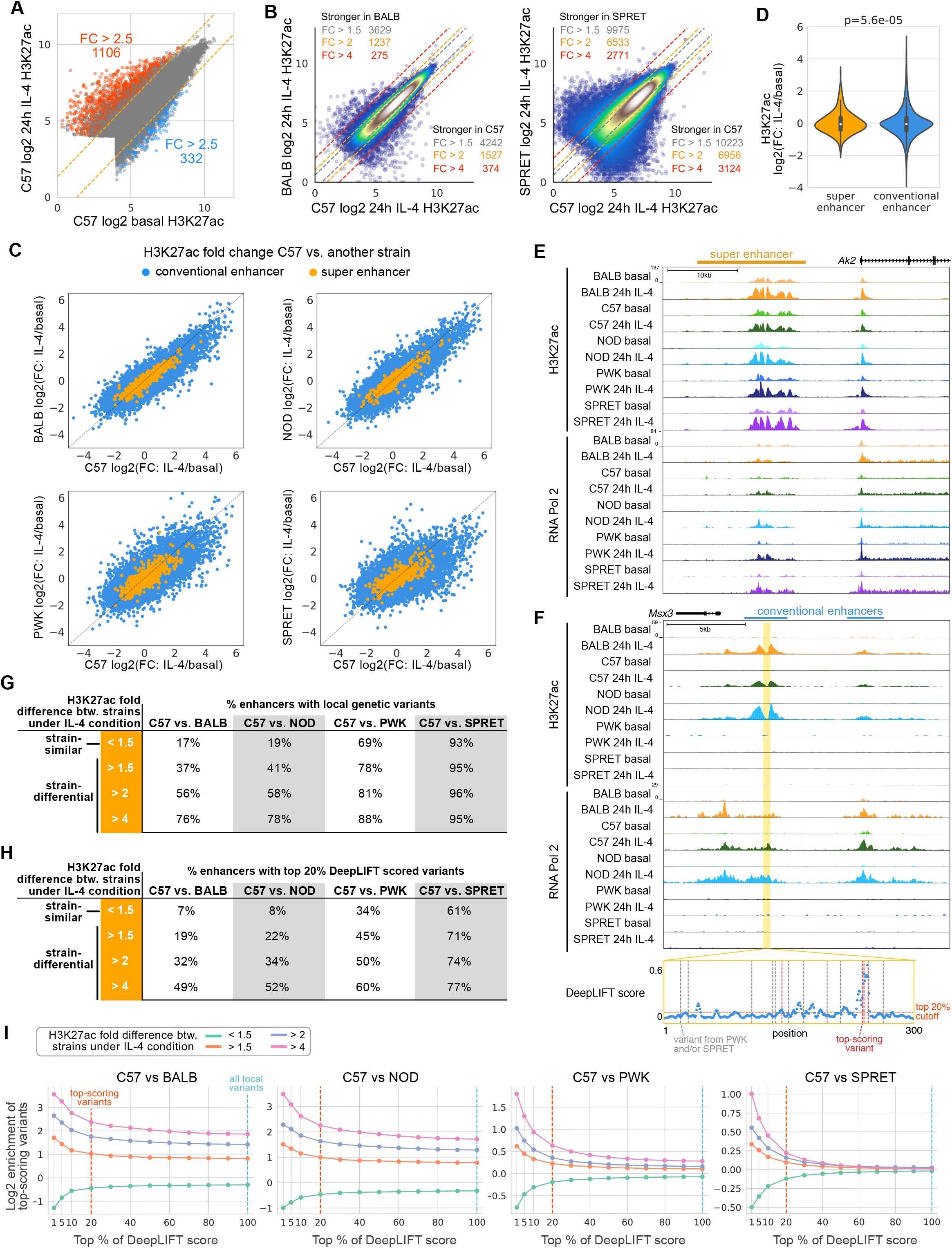
Divergent IL-4 response is associated with strain-differential IL-4 enhancer activation. A. Log2 H3K27ac signal at ATAC peaks in C57 BMDMs under basal and IL-4 conditions. B. Comparison of H3K27ac signal between C57 and BALB or SPRET under the 24h IL-4 condition. C. Log2 H3K27ac fold changes after 24h IL-4 in C57 versus other strain in enhancers. D. Distributions of IL-4 H3K27ac log2 fold changes, Levene’s test was performed to test response differences in conventional versus super enhancers. E. *Ak2* super enhancer responsive to IL-4 and conserved across all strains. F. *Msx3* IL-4 induced enhancer in C57, BALB and NOD, but not PWK and SPRET BMDMs. Absolute DeepLIFT scores indicate predicted importance of single nucleotides for enhancer activity. Dotted lines represent locations of PWK or SPRET variants. G-H. Enhancers were categorized into strain-similar and strain-differential based on fold differences in H3K27ac between C57 and one of the other strains. Table with percentages of enhancers containing local genetic variants in G and the percentage of enhancers that contain predicted functional variants in H. I. Log2-scaled enrichment of enhancers with variants at top-scoring positions based on DeepLIFT scores. The enrichment was calculated by (% enhancers in one category with top variants)/(% all enhancers with top variants). 2G and 2H are based on the top 100% and 20%, respectively.

We next compared the fractions of enhancers containing variants in strain-similar enhancers (<1.5-fold differences in H3K27ac between strains) to strain-differential enhancers at increasing levels of difference (fold differences >1.5 to >4; Figure 2G). The fraction of enhancers containing variants at strain-similar enhancers ranged from 17-20% in the strains most similar to C57 (BALB and NOD) to 69-93% in the most genetically divergent strains (PWK and SPRET). As expected, the fraction of enhancers containing variants increased with increasing levels of difference, except for SPRET which may have reached a saturation of variation capacity (Figure 2G). These findings are consistent with local variants affecting enhancer activity, but also indicate that a substantial fraction of even strongly strain-differential IL-4 induced enhancers lack such variants, consistent with previous findings for strain-specific enhancers overall (Link et al., 2018a).

In an effort to distinguish silent variants from those affecting enhancer activity, we trained a DeepSEA convolutional neural network to classify enhancers as active or inactive under the 24h IL-4 condition based on local sequence context (Zhou and Troyanskaya, 2015). The training data consisted of enhancers active under IL-4 conditions (positive data) and random background (negative data). The area under the receiver operating characteristic curve (auROC) was 0.894 on test data. We then used DeepLIFT (Shrikumar et al., 2017) to compute the importance score of each nucleotide based on the model’s classification decision. Variants at positions with top importance scores within surrounding 300-bp enhancer regions are hypothesized to affect enhancer activity. We considered variants residing in the top 20% of importance scores for each region as predicted functional variants. The *Msx3* enhancer in Figure 2F illustrates four predicted functional variants out of fourteen variants in PWK and SPRET (red dotted lines). By focusing on top-scoring variants rather than all local variants, we saw an expected overall decreased percentage of enhancers with top-scoring variants (Figure 2H, S2H). On the other hand, enrichment of predicted functional variants increases as a function of importance score threshold and is strongest for enhancers that show the highest differences across strains (Figure 2I). This is true when considering all strains, including SPRET. These results reveal a quantitative impact of variants affecting enhancer under IL-4 treatment conditions and suggest the extent to which a deep learning approach can distinguish potentially functional variants from the silent variants.

### IL-4 activated enhancers use existent promoter-enhancer interactions to regulate gene activity

Interpretation of effects of genetic variation on distal regulatory elements is facilitated by knowledge of cell-specific enhancer-promoter interactions (Nott et al., 2019). To identify connections of IL-4-responsive enhancers to target promoters, we performed HiChIP using an antibody to H3K4me3 (Mumbach et al., 2016) in C57 BMDMs under basal conditions and after 24h of IL-4 treatment. HiChIP interactions are exemplified in Figure 3A at the *Slc7a2* locus, a gene that becomes maximally activated after 24h of IL-4 treatment (Figure 3B) and connects primarily to an enhancer-like region within the *Mtmr7* gene which itself is expressed at negligible levels (Figure 3B). Although we observed instances of IL-4-specific interactions (e.g. yellow loops), a differential interaction analysis was unable to identify significantly different interactions (Figure S3A). However, IL-4 activated promoters mostly interact with IL-4 activated enhancers (Fisher’s exact test, p=2.2E-16) and repressed promoters strongly interact with IL-4 repressed enhancers (p=1.2E-15, Figure 3C).

**Figure 3.**
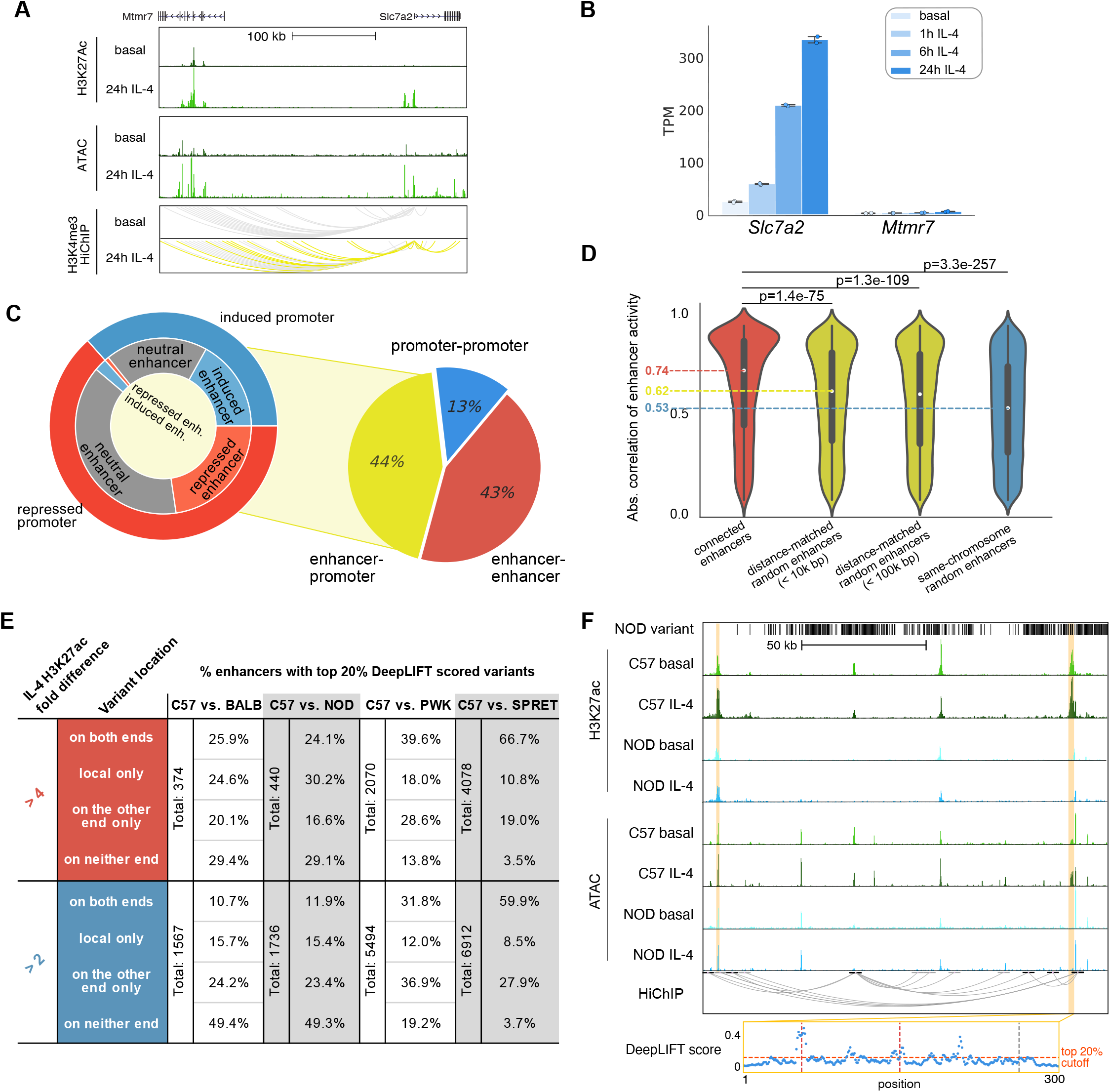
IL-4 enhancers use existent promoter-enhancer interactions to regulate gene activity. A. HiChIP indicates the *Slc7a2* promoter is highly connected with several IL-4 activated enhancers. B. *Slc7a2* and *Mtmr7* gene expression upon IL-4 stimulation. C. HiChIP interactions. Interactions between promoters and enhancers (right), and enhancer-promoter connections overlapping with IL-4 responsive regulatory elements in C57 BMDMs (left). Outer ring indicates induced or repressed promoters, while inner ring indicates their connected enhancers associated with IL-4-induced, IL-4 repressed or IL-4 neutral H3K27ac. D. Correlations of H3K27ac signal between connected enhancers compared to other groups of enhancers using Mann–Whitney U test. E. Table representing enhancers containing genetic variants locally or at connected elements in pairwise comparisons between C57 and other strains. F. Strain-differential enhancer between C57 and NOD where genetic variants were absent locally but present at a connected enhancer with two DeepLIFT predicted functional variants (red dotted lines).

Although the HiChIP assay is designed to capture promoter-enhancer interactions based on preferential occurrence of H3K4me3 at promoters, we also recovered 242,837 pairs of interactive enhancers (Figure 3C), consistent with more than one enhancer being in local proximity of a target promoter. To investigate whether the interactive enhancers are functionally related, we computed the correlation of H3K27ac signal within each enhancer pair across the five strains. The correlations between interactive enhancers were significantly stronger than those between non-interactive enhancers (Figure 3D, S3B). Based on this result, we investigated whether enhancer-enhancer interactions could explain strain-differential enhancers lacking top-scoring variants with potential influence on enhancer activities (Figure 2H). Among 374 interactive enhancers exhibiting a >4-fold difference in H3K27ac signal between BALB and C57 under the IL-4 condition, the original ~50% of strain-differential enhancers with top 20% predicted functional variants was further split into 25.9% that had top-scoring variants on both ends and 24.6% that had only local top-scoring variants (Figure 3E upper left). For the PWK and SPRET comparisons, a larger proportion of enhancers had top-scoring variants on both ends, consistent with their higher percentages for local top-scoring variants (Figure 2H). An additional 16.6%-28.6% of strain-differential enhancers, despite a lack of local top-scoring variants, were connected to enhancers containing top-scoring variants that potentially affected their activity (Fisher’s exact test p=4E-33 for BALB, 7E-27 for NOD, 2E-7 for PWK, 0.06 for SPRET, compared to strain-similar enhancers). The marginal significance of SPRET compared to high significance in all other strain comparisons is driven by the high frequency of variants in strain-similar enhancers. Reducing the fold change requirement to 2-fold yielded a smaller proportion of strain-differential enhancers containing local variants overall but significantly increased the proportion having top-scoring variants on the connected ends only (Fisher’s exact test p=0.05 for BALB, 0.001 for NOD, 5E-12 for PWK, 2E-26 for SPRET), suggesting that local variants have a stronger effect on inducing differential activation than variants at connected enhancers (Figure 3E lower panels, S3C). The reciprocal relationships of p values for these two comparisons are driven by the frequencies of variants in strain-similar enhancers. Figure 3F illustrates an enhancer affected by genetic variants at the connected enhancer. The enhancer highlighted on the left is significantly more activated in C57 than NOD. This region lacks local variants in NOD but is connected to another enhancer ~100 kb away containing multiple variants that are predicted to affect activity (highlighted on the right). These findings are consistent with genetic variants at an enhancer influencing the activity states of other enhancers that lack local functional variants within the same connected network (Grubert et al., 2015; Waszak et al., 2015).

### Motif mutation analysis identifies motifs that are functionally associated with IL-4 induced enhancer activity

IL-4 rapidly activates a set of enhancers, the majority of which exhibit maximal H3K27ac at 1h - 6h that returns to (near) basal levels by 24h (Figure 4A, top three clusters) when most gene expression changes were found (Figure S1B). Others are long-lasting or become activated at later timepoints (Figure 4A, bottom three clusters). *De novo* motif enrichment analysis of enhancers exhibiting >2.5-fold increase in H3K27ac and RNA Pol2 at 1h, 6h and 24h (Figure S2A) recovered a STAT6 motif as the most enriched motif for all timepoints (Figure 4B). Motifs for the lineage determining factors PU.1 and AP-1 family members were also recovered in all three classes of enhancers. Notably, an EGR2 motif was significantly enriched among enhancers induced at 24 hours.

**Figure 4.**
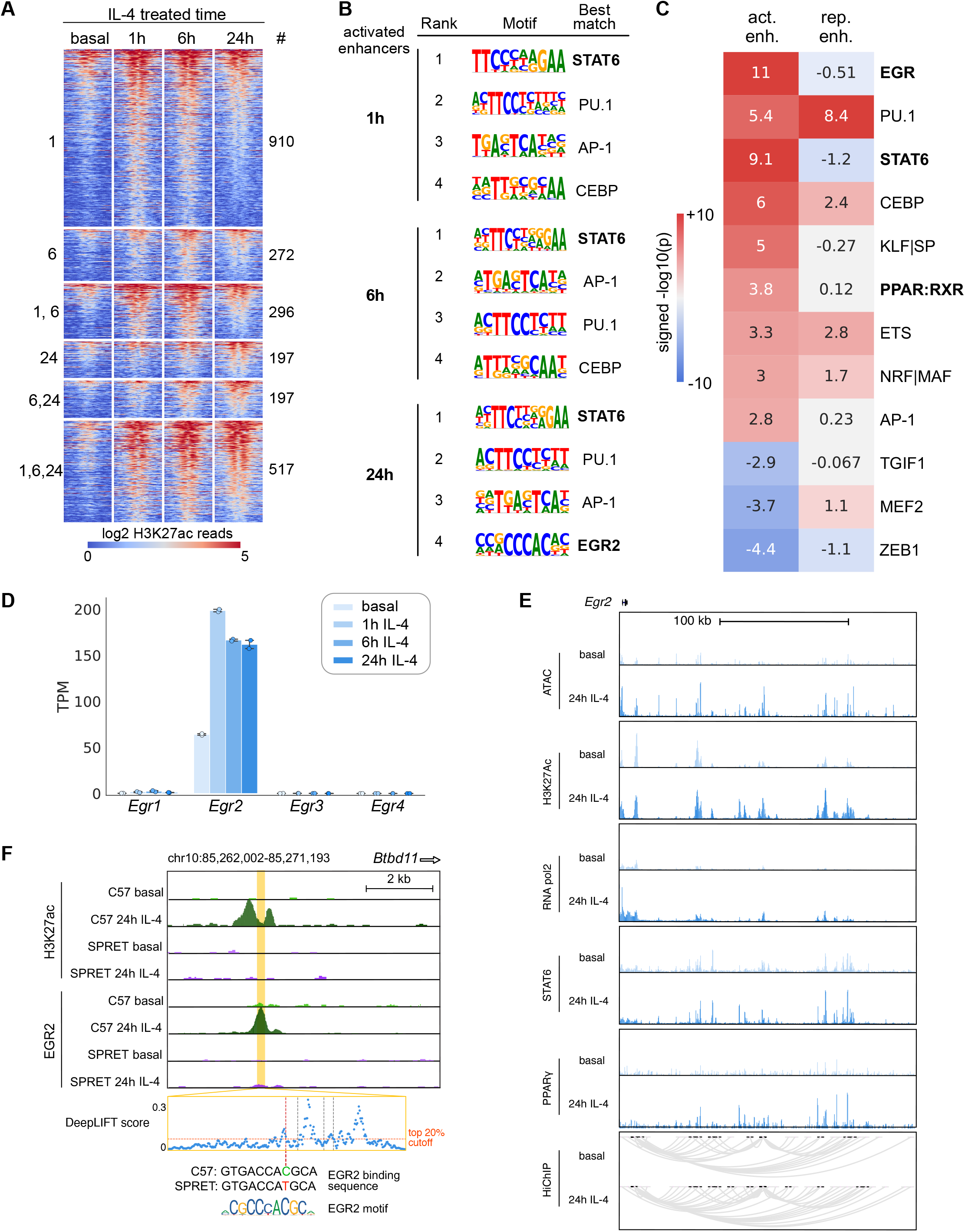
Motif mutation analysis identifies motifs functionally associated with IL-4 induced enhancer activity. A. Heatmap showing the effects of 1h, 6h and 24h IL-4 stimulation on enhancer activation based on H3K27ac abundance. B. Top motifs enriched at ATAC-seq peaks exhibiting gained H3K27ac at 1, 6, or 24 hours IL-4. C. MAGGIE motif mutation analysis on strain-differential activated and repressed enhancers after 24h IL-4. D. *Egr* gene expression in C57 BMDMs under basal conditions and after stimulation with IL-4, ****q<0.0001, compared to basal. E. *Egr2* promoter connected to several upstream enhancers in C57 BMDMs as determined by H3K4me3 HiChIP. Connected enhancers bound by SDTFs STAT6 and PPARg display increased H3K27ac and RNA Pol2 by IL-4. F. Example of a strain-differential activated enhancer upstream of the *Bdb11* gene based on IL-4-induced H3K27ac signal in C57 but not in SPRET BMDMs, which is supported by binding of EGR2 and a functional variant predicted by DeepLIFT that mutates the EGR2 motif.

As a genetic approach to identify functional transcriptional factor binding motifs, we assessed the quantitative impact of the genetic variation provided by the five different strains of mice on the IL-4 response of enhancers using the motif mutation analysis tool MAGGIE (Shen et al., 2020). This analysis identified more than a dozen motif clusters in which motif mutations were significantly associated with strain-differential IL-4 activated or repressed enhancers (Figure 4C, S4A). The EGR motif was found as the top motif associated with enhancer activation at the 24h treatment time, as well as motifs of known SDTFs STAT6 and PPARg and macrophage LDTFs PU.1, AP-1 and CEBP (Figure 4C). We also found KLF motifs associated with IL-4 enhancer activation, which fits with increased KLF4 expression by IL-4 (Figure S4B), and an NRF motif associated with both enhancer activation and repression (Figure 4C).

The identification of STAT6 and PPARg motif mutations as being functionally associated with strain-differential IL-4 activation is consistent with substantial prior work demonstrating the importance of these factors in regulating IL-4-dependent gene expression (Czimmerer et al., 2018; Daniel et al., 2018). Out of the Early Growth Response (EGR) family members only *Egr2* is expressed in unstimulated BMDMs and rapidly induced after IL-4 stimulation (Figure 4D, S4C). *Egr2* has also been associated with late IL-4 enhancer activation in a recent study (Daniel et al., 2020). Examination of the *Egr2* locus indicates IL-4 induced binding of STAT6 and PPARg to a set of upstream super enhancers that gain H3K27ac and RNA Pol2 signal after IL-4 stimulation (Figure 4E). These super enhancers were observed in BMDMs of all five different strains (Figure S4D) that are strongly connected to the *Egr2* promoter in C57 BMDMs as indicated by H3K4me3 HiChIP interactions. Overall, the induction of *Egr2* by IL-4 and the effect of EGR motif mutations on the activity states of IL-4-induced enhancers suggests a functionally important role of EGR2 in contributing to IL-4 induced gene expression in BMDMs.

### IL-4 induced EGR2 contributes to late IL-4 enhancer activation

To investigate whether EGR2 activates IL-4 dependent enhancers, we performed ChIP-seq for EGR2 under basal and 24h IL-4 treatment conditions. This confirmed the prediction that mutations in EGR binding sites contribute to strain-differential enhancer activation by altering the binding of EGR2. An example is provided by the *Btbd11* enhancer, which is IL-4 inducible in C57, but not in SPRET BMDMs (Figure 4F). Consistent with a near complete loss of EGR2 binding in SPRET, a C-to-T variant in SPRET mutated an EGR2 motif and was predicted as functional by DeepLIFT.

Relative binding of EGR2 in C57 BMDMs as a function of time following IL-4 treatment is represented in Figure 5A. Overall, IL-4 treatment resulted in a marked expansion of the EGR2 cistrome after stimulation with IL-4 (Figure 5B). In addition to an increase in the number of EGR2 peaks after IL-4, the peak intensity also increased (Figure 5A). Accordingly, most EGR2 peak intensities reached maximum values at 6 and 24h, while STAT6 binding is high immediately after 1h already and then slowly decreases in peak intensity (Figure S5A). The IL-4 induced EGR2 peaks were associated with increased H3K27Ac and RNA Pol2 signal, consistent with a role of EGR2 in late enhancer activation (Figure 5C, S5B).

**Figure 5.**
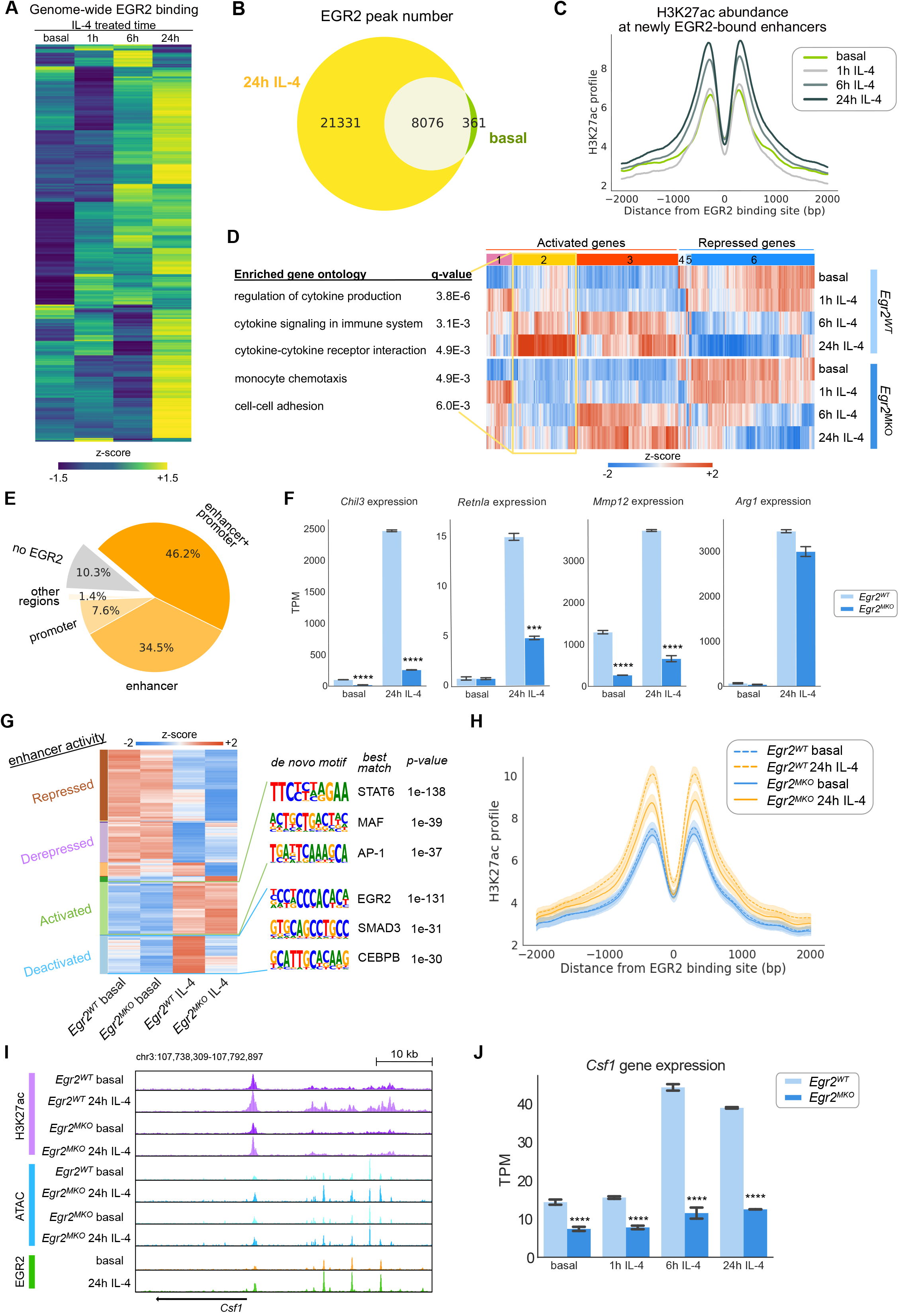
IL-4 induced EGR2 contributes to late IL-4 enhancer activation. A. Heatmap displaying EGR2 ChIP-seq binding intensity after IL-4 stimulation over time in C57 BMDMs. B. EGR2 binding sites after 24h IL-4 compared to the basal in C57 BMDMs. C. H3K27ac profiles at 24h IL-4 induced intergenic and intronic EGR2 peaks in C57 BMDMs. D. Expression of IL-4 regulated genes in *Egr2^WT^* and *Egr2^MKO^* BMDMs, top GO terms displayed for cluster 2. E. EGR2 binding to promoters and enhancers of EGR2-dependent IL-4 target genes. F. *Mmp12, Retnla* and *Chil3* genes affected by and *Arg1* not affected by *Egr2* deletion, ***q<0.001, ****q<0.0001, compared to *Egr2^WT^.* G. Enhancer activity of IL-4 regulated enhancers in *Egr2^WT^* and *Egr2^MKO^* BMDMs. Motifs at EGR2-dependent and -independent IL-4 induced enhancers enriched compared to the other group. H. H3K27ac profiles at IL-4 induced EGR2 binding sites in *Egr2^WT^* and *Egr2^MKO^* BMDMs. 90% confidence intervals are shown together with the average profiles. I. EGR2-bound *Csf1* enhancers after 24h IL-4 stimulation. J. *Csf1* gene expression after IL-4 stimulation in *Egr2^WT^* and *Egr2^MKO^* BMDMs, ****q< 0.0001.

To extend these analyses, we crossed *Egr2^fl/fl^* (*Egr2^WT^,* Du et al., 2014) with *LyzM*-Cre^+^ mice to obtain *LyzM*-Cre^+^ *Egr2*^flfl^ (*Egr2*^MKO^) mice. This resulted in efficient deletion of *Egr2* in BMDMs (Figure S5C-D). RNA-seq data from *Egr2^WT^* and *Egr2^MKO^* BMDMs indicated that at the mRNA level, EGR2 is regulating ~40% of the 24h IL-4 induced genes (Figure 5D). *Egr2* deletion mainly affects late IL-4 targets genes (yellow cluster, #2) at 6 and 24 hours. Early IL-4 induced genes at 1h and 6h in the purple cluster (#1) are not affected by *Egr2* deletion (Figure 5D, 54E-F, Table S3). GO analysis shows that pathways regulating cytokine production, monocyte chemotaxis and cell-cell adhesions are among the top significant terms that are downregulated in *Egr2^MKO^* BMDMs (Figure 5D). The majority of these EGR2 targets genes have EGR2 binding in their promoter and/or enhancer (Figure 5E), which is consistent with the effects of deletion being direct consequences of EGR2 loss at gene regulatory elements. Examples of EGR2-dependent IL-4 target genes are typical IL-4 response marker genes *Mmp12, Retnla* (Fizz1) and *Chil3* (Ym1). In contrast, gene expression of another classical IL-4 responsive gene *Arg1* is not affected by deletion of *Egr2* (Figure 5F).

Next, we investigated the effects of *Egr2* deletion on enhancer activity. We found that ~40% of the 24h IL-4 induced enhancer activity was significantly decreased in *Egr2^MKO^* BMDMs (blue cluster, Figure 5G, Figure S5G-I). In concordance, the IL-4 induction in H3K27ac was not observed at IL-4 induced EGR2 binding sites in *Egr2^MKO^* BMDMs (Figure 5H). When performing motif analysis of EGR2-dependent IL-4 activated enhancers, we found EGR2 as the most significantly enriched motif in this cluster with the EGR2-independent cluster as background and the SMAD3 and CEBP motifs as second and third hits (Figure 5H). In the EGR2-independent cluster, the STAT6 motif was most significantly enriched (Figure 5G), with EGR2-dependent cluster as a background. A clear example of an IL-4 inducible and EGR2-dependent enhancer is the *Csf1* enhancer where IL-4 increased EGR2 binding and H3K27 acetylation (Figure 5I). As a result, gene expression increased after 6 and 24 hours of IL-4 treatment in *Egr2^WT^* BMDMs but maintained at a low level in *Egr2*^MKO^ BMDMs (Figure 5J).

### Collaborative and hierarchical transcription factor interactions at IL-4 dependent enhancers

Analysis of the genome-wide binding patterns of EGR2, STAT6 and PPARg indicated intensive cobinding at IL-4 activated enhancers (Figure S6A). To achieve highly strain-differential responses to IL-4 that are observed at the level of gene expression (Figure 1) and enhancer activation (Figure 2), these factors are hypothesized to exert their transcriptional effects via correspondingly divergent genomic binding patterns. ChIP-seq analysis of EGR2, STAT6, PPARg, C/EBPβ and PU.1 in each strain under basal and 24h IL-4 conditions confirmed this. For example, over 4,000 EGR2 binding sites exhibited more than 4-fold differences in normalized tag counts between C57 and BALB, and almost 9,000 between C57 and SPRET (Figure 6A). Similar relationships are observed for STAT6 (Figure 6B) and each of the other factors (not shown).

**Figure 6.**
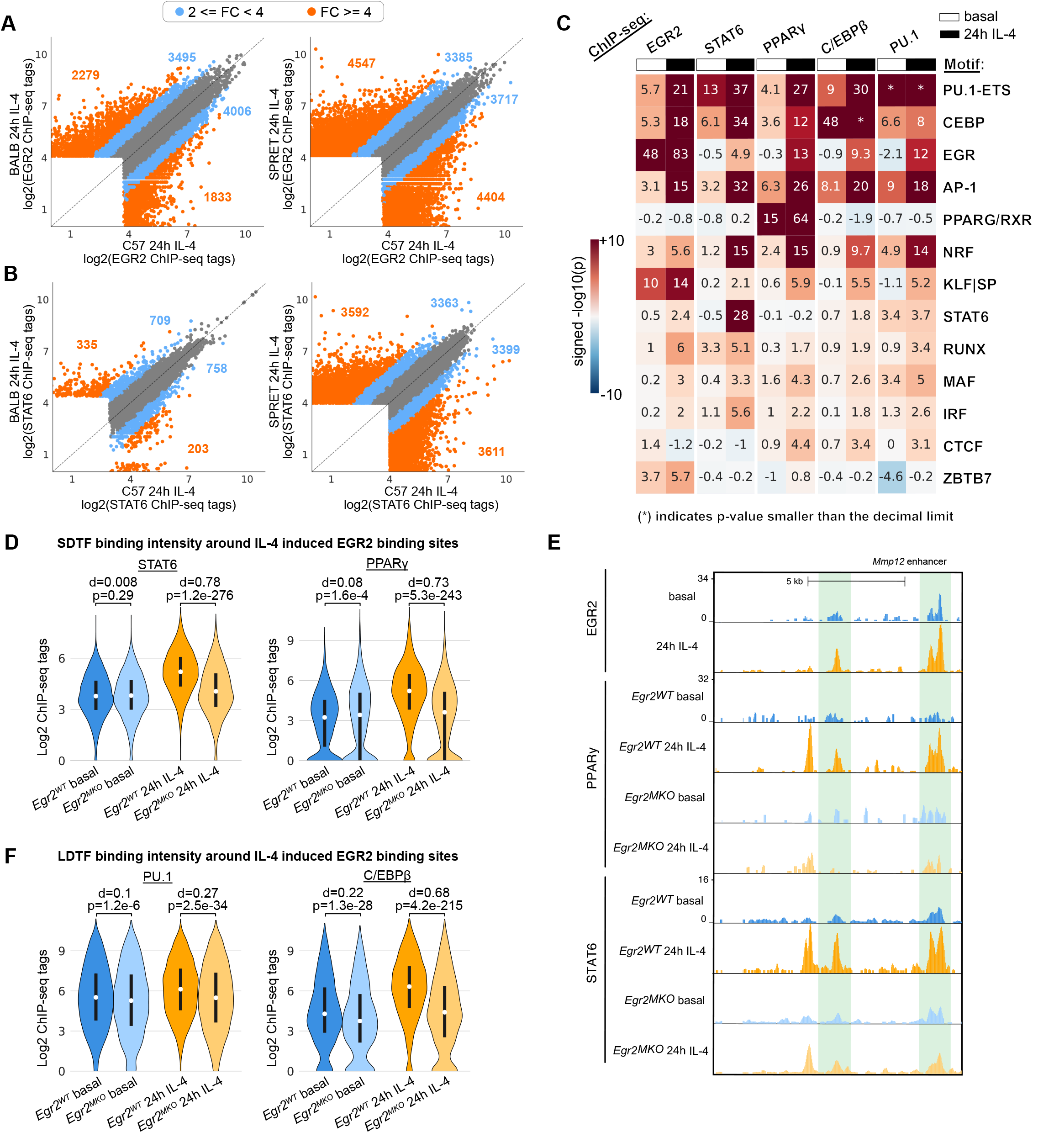
Collaborative and hierarchical transcription factor interactions at IL-4 enhancers. A-B. Scatter plots comparing binding of EGR2 (A) and STAT6 (B) in C57 versus BALB and C57 versus SPRET IL-4 stimulated BMDMs. C. Functional motifs from MAGGIE analysis at EGR2, STAT6, PPARg, C/EBPβ and PU.1 peaks. D. STAT6 and PPARg binding at IL-4 induced EGR2 peaks in *Egr2^WT^*and *Egr2^MKO^* BMDMs. Cohen’s d effect size and p-values from Mann–Whitney U tests are shown. E. Co-binding of STAT6, EGR2, and PPARg at the *Mmp12* enhancer. F. C/EBPβ and PU.1 binding at IL-4 induced EGR2 peaks in *Egr2^WT^*and *Egr2^MKO^* BMDMs.

The strain-differential binding patterns of SDTFs and LDTFs enabled motif mutation analysis to study the importance of motifs for SDTF and LDTF binding (Figure 6C). As a validation, LDTF and SDTF binding depended on their own motifs (e.g., PU.1 motif mutation was significantly associated with PU.1 binding, indicating that when PU.1 binding is lost in one strain, it is often found that the PU.1 motif score is reduced in that strain compared to the other). In addition, mutations in the motifs of LDTFs PU.1, C/EBP and AP-1 influence the binding of all LDTFs and SDTFs, which fits with earlier observations (Heinz et al., 2013). We found that the PPAR motif is only significant for PPARg binding, and likewise, the STAT6 motif is not associated with binding of other SDTFs or LDTFs but STAT6. Interestingly, we found that mutations in EGR2 motifs are significantly associated with binding of SDTFs STAT6 and PPARg and LDTFs PU.1, C/EBPβ under

IL-4 conditions, but not under basal conditions. These analyses also provided evidence for functional roles of several additional transcription factors. Mutations in NRF motifs were strongly associated with the IL-4-dependent binding of all SDTFs and LDTFs. Both NRF1 and NRF2 are expressed in BMDMs (Figure S4B) and are involved in lipid metabolism and stress responses (Eichenfield et al., 2016; Kobayashi et al., 2016; Widenmaier et al., 2017). Mutations in KLF motifs were strongly associated with EGR2 binding under both basal and IL-4 conditions. KLF2, KLF4 and KLF6 are expressed in BMDMs (Figure S4B) and KLF4 has previously been associated with antiinflammatory roles in macrophages (Liao et al., 2011). Mutations in IRF motifs were moderately associated with IL-4-dependent STAT6 binding. Multiple IRFs, including IRF4, are expressed in BMDMs (Figure S6B) and IRF4 has previously been linked to macrophage polarization by IL-4 (El Chartouni et al., 2010; Satoh et al., 2010).

A prediction emerging from the analysis results above is that EGR2 should have a small effect on the co-binding of SDTFs and LDTFs under basal conditions and a significant effect following 24h of IL-4 treatment. To examine this prediction, we performed ChIP-seq for STAT6, PPARg, PU.1 and C/EBPβ in *Egr2^WT^* and *Egr2^MKO^* BMDMs and evaluated their binding in the vicinity of IL-4 induced EGR2 binding sites. Deletion of *Egr2* had little effect on PPARg and STAT6 binding under basal conditions and a much greater effect following 24h of IL-4 treatment (Figure 6D). As an example, in *Egr2^MKO^* BMDMs, PPARg and STAT6 binding was found decreased at the *Mmp12* enhancer at sites where EGR2 normally binds (Figure 6E). Similarly, PU.1 and C/EBPβ binding was more significantly affected by *Egr2* deletion under the IL-4 condition than the basal condition (Figure 6F). In concert, these findings provide evidence for collaborative and hierarchical interactions between PU.1, C/EBPs, AP-1, STAT6, PPARg and EGR2 as major drivers of late enhancer activation in response to IL-4.

### Determinants of absolute levels and dynamic responses of IL-4 responsive enhancers

We next investigated the possibility that the mutational status of the dominant motifs recovered by MAGGIE analysis was sufficient to predict qualitative patterns of strain-differential responses of IL-4 induced enhancers. Following the classification of strain-differential mRNA responses (Figure 1), we used H3K27ac to define three different categories of strain-differential IL-4-induced enhancers (Figure 7A, left column): enhancers exhibiting lower levels of basal activity in the lowly induced strain (**low basal**); enhancers with a similar level of basal activity (**equal basal**); enhancers in which a lack of IL-4 induced activity was associated with relatively higher basal activity compared to the more responsive strain (**high basal**). Using these criteria, we identified 760 low basal, 2797 equal basal and 2013 high basal enhancers from all pairwise comparisons of the five strains that exhibited >2-fold differences in H3K27ac induction (Figure 7B). Low basal, equal basal and high basal enhancers are exemplified by enhancers associated with the *Treml2, Ripk2* and *Cd36* genes, respectively (Figure 7C-E, S7A-C).

**Figure 7.**
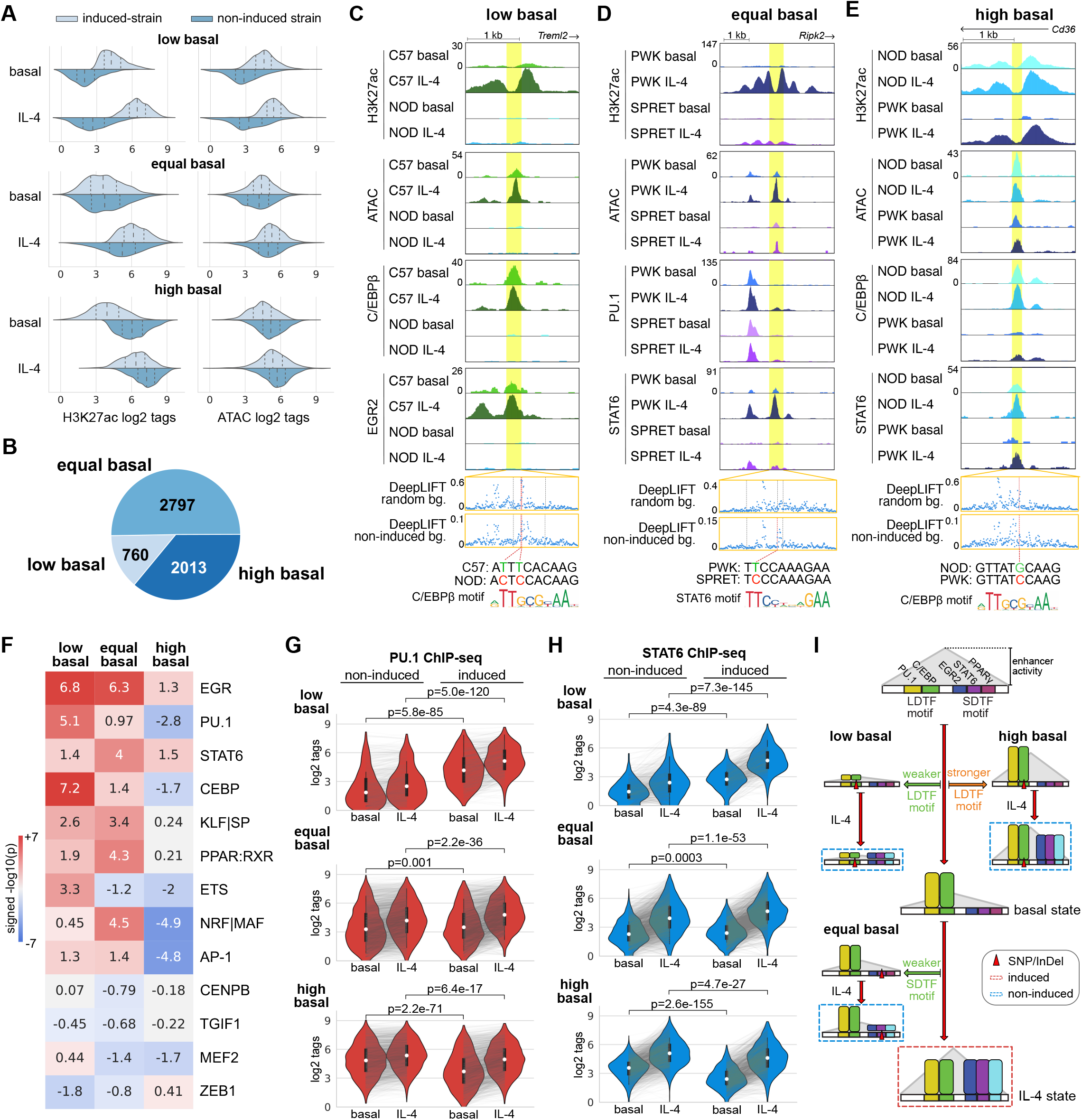
Determinants of absolute levels and dynamic responses of IL-4 enhancers. A. Three different categories of strain-differential IL-4 activated enhancers with distributions of ATAC and H3K27ac signal. Dashed lines in each distribution indicate quartiles. B. Numbers of enhancers in the three categories. C-E. Example of low (C), equal (D) and high (E) basal enhancers with high impact variants predicted by DeepLIFT. F. MAGGIE motif mutation analysis on different categories of enhancers. G-H. Binding intensities of PU.1 (G) and STAT6 (H) in non-induced and induced strains at different categories of enhancers. I. Graphical representation of the general mechanisms for different categories of IL-4 induced enhancers.

Consideration of chromatin accessibility as determined by ATAC-seq further uncovered potential mechanisms that distinguished the three enhancer categories (Figure 7A, right column). The enhancers in the low basal category showed low to absent basal ATAC signal in non-induced strains, suggesting a lack of LDTFs under the basal condition to pre-occupy chromatin required for subsequent recruitment of SDTFs after IL-4 stimulation. In contrast, high basal enhancers exhibited a higher basal level of ATAC in non-induced strains compared to induced strains (Figure 7A, right column), suggesting stronger LDTF binding in non-induced strains under the basal condition. Different from the other categories, equal basal enhancers exhibited similar levels of chromatin accessibility under both basal and IL-4 conditions between comparative strains, suggesting that the recruitment of SDTFs might be the key determinant for the strain difference instead of basal LDTF binding.

To test the hypotheses above regarding the different determinants for the three categories of enhancers, we performed MAGGIE motif mutation analysis on each category of enhancers. We found that mutations in motifs of LDTFs PU.1/ETS and C/EBP were associated with low basal enhancers and resulted in better motifs in induced strains, while mutations in motifs of SDTFs EGR, STAT6, PPAR and NRF/MAF were associated with the equal basal category leading to better motifs in induced strains (Figure 7F, S7D). Mutations in EGR motifs were also associated with the low basal category, suggesting another role of EGR2 as a pioneering factor under the IL-4 condition, supported by the significant decrease in open chromatin under IL-4 conditions after deletion of *Egr2* (Figure S5G). Of particular interest, the high basal category of enhancers was most strongly associated with negative significance scores for LDTF PU.1, C/EBP and AP-1 as well as NRF/MAF, meaning better motifs in non-induced strains (Figure 7F).

We validated these findings with our ChIP-seq data by examining the binding profiles of PU.1, C/EBPβ, STAT6, PPARg and EGR2 in three categories of enhancers. In low basal enhancers, we saw significantly reduced binding of PU.1 and C/EBPβ in non-inducible strains under both basal and IL-4 conditions (Figure 7G, S7E). This pattern was accompanied by significantly weaker binding of SDTFs STAT6, EGR2 and PPARg after IL-4 stimulation (Figure 7H, S7F). The example in Figure 7C showed the absence of C/EBPβ binding in NOD under the basal condition likely due to two local variants at high-scored positions according to DeepLIFT that together mutated a C/EBP motif. Upon IL-4 stimulation, neither C/EBPβ nor EGR2 was further recruited. For equal basal enhancers, we found that PU.1 and C/EBPβ binding was similar under basal conditions in induced and non-induced strains (Figure 7G, S7E). Upon IL-4 stimulation, the induced strains displayed significantly stronger binding of SDTFs STAT6, EGR2 and PPARg (Figure 7H, S7F). In the example in Figure 7D, STAT6 binding was strongly induced by IL-4 at the *Ripk2* enhancer in PWK but was absent in SPRET. Despite the clear difference in STAT6 binding, none of the local variants between the two strains was predicted functional when using a neural network model trained with random genomic backgrounds. To better capture the sequence patterns relevant for enhancer activation, we retrained neural networks using non-induced enhancers as the background, which emphasized a relatively divergent set of k-mers, especially those matched with SDTF motifs (Figure S7G). As a result, our retrained model assigned a high DeepLIFT score to one of the nucleotides in a STAT6 motif that was mutated by a variant in SPRET (Figure 7D). For high basal enhancers, we found stronger binding of not only the LDTFs PU.1 and C/EBPβ (Figure 7G, S7E) but also the SDTFs STAT6 and PPARg (Figure 7H, S7F) in non-induced strains under basal conditions. For example, high basal levels of C/EBPβ and STAT6 binding were observed at the *Cd36* enhancer in NOD mice (Figure 7E). The only local variant in PWK was at a predicted functional position and mutated a C/EBP motif likely causing the low basal C/EBPβ binding in PWK. In concert, these analyses validated the importance of LDTF motif mutations as primary determinants of differential enhancer activation in low basal and high basal enhancers, while also demonstrating the expected consequences of SDTF motif mutations in determining strain-differential activation of equal basal enhancers (Figure 7I).

## Discussion

Here, we report a systematic investigation of the effects of natural genetic variation on signal-dependent gene expression by exploiting the highly divergent responses of BMDMs from diverse strains of mice to IL-4. Unexpectedly, despite broad conservation of IL-4 signaling pathways and downstream transcription factors in all five strains, only 26 of more than 600 genes observed to be induced >2-fold by IL-4 at 24 hours reached that level of activation in all five strains and more than half were induced in only a single strain. To the extent that this remarkable degree of variation observed in BMDMs occurs in tissue macrophages and other cell types *in vivo,* it is likely to have significant phenotypic consequences with respect to innate and adaptive immunity, tissue homeostasis and wound repair. Notably, only ~25% of the variation in response to IL-4 was due to altered dynamic ranges in the context of an equivalent level of basal expression. Nearly half of the genes showing strain-specific impairment in IL-4 responsiveness exhibited low basal activity, whereas lack of induction was associated with constitutively high basal levels of expression in the remaining ~25%. These qualitatively different patterns of strain responses to IL-4 imply distinct molecular mechanisms by which genetic variation exerts these effects.

Motif mutation analysis of strain-differential enhancer activation recovered a dominant set of motifs recognized by known LDTFs PU.1, C/EBPβ and AP-1 family members, as well as motifs recognized by SDTFs STAT6 and PPARg that have been previously established to play essential roles in the IL-4 response. In addition, effects of mutations in motifs for EGR, NRF and KLF also strongly implicate these factors as playing important roles in establishing basal and induced activities of IL-4 responsive enhancers, which was genetically confirmed for EGR2. It will be of interest in the future to perform analogous studies of NRF and KLF factors.

Analysis of strain-differentially activated enhancers revealed qualitative differences in basal and IL-4-dependent activity that were analogous to the qualitative differences observed for strain-differentially activated genes. As expected, sequence variants reducing the affinity of SDTFs STAT6, PPARg and EGR2 were the major forms of variation resulting in strain-differential IL-4 induction of equal basal enhancers. From the standpoint of interpreting the effects of non-coding variation, these types of sequence variants are silent in the absence of IL-4 stimulation. As also expected, sequence variants strongly reducing the binding affinity of LDTFs prevented the generation of open chromatin required for subsequent binding of SDTFs. Such variants are thus expected to result in loss of enhancer function in a signal-independent manner. Of particular significance, these analyses also provide strong evidence that quantitative variation in suboptimal motif scores for LDTFs is a major determinant of differences in the absolute levels and dynamic range of high basal enhancers across strains. The importance of low affinity motifs in establishing appropriate quantitative levels of gene expression within a given cell type and cell specificity across tissues has been extensively evaluated (Crocker et al., 2015; Farley et al., 2015; Kribelbauer et al., 2019). Here we present evidence that improvement of low affinity motifs for LDTFs not only increases basal binding of the corresponding transcription factor but is also associated with increased basal binding of STAT6 and PPARg, thereby rendering their actions partially or fully IL-4 independent. These findings thus provide evidence that quantitative effects of genetic variation on LDTF motif scores play major roles in establishing different absolute enhancer activity levels and dynamic ranges of their responses to IL-4 that are observed between strains.

To go beyond the discovery of mechanisms mediating the IL-4 response using natural genetic variation, a major objective of these studies was to use the resulting data sets as the basis for interpreting and predicting the effects of specific variants. As expected, enhancers exhibiting strain specific differences in IL-4 responses were significantly enriched for sequence variants. However, the background frequencies of variants in the much larger sets of strain-similar enhancers ranged from 17% to 93%, consistent with the vast majority of such variants being silent and underscoring the challenges of discriminating them from functional variants. The application of recently developed deep learning approaches illustrates both the potential of these methods to improve predictive power as well as their current limitations. Nucleotides predicted by DeepLIFT to be of functional importance frequently intersected with variants at strain-differential enhancers that significantly altered LDTF or SDTF motifs, with over 8-fold enrichment in enhancers with strongest strain differences (top 1% variants for C57 vs. BALB comparison, Fig. 2I), strongly suggesting causality. Even though DeepLIFT scored a significant fraction of variants present in strain-similar enhancers with low importance, a large fraction of remaining strain-similar enhancers contained variants associated with high DeepLIFT scores, most likely representing false positives. Further, we found that the highest scoring variants in some cases depended on the choice of data used to train the convolutional neural network (e.g. using random vs. non-induced enhancers as negative training examples). This observation has significant implications with respect to application of deep learning models to identify potential functional variants in disease contexts. The data sets generated by these studies will therefore provide an important resource for further improvements in methods for interpretation of local genetic variation.

These analyses further indicated that 20%-50% of the most divergent IL-4-responsive enhancers lacked any functional variants in the proximity of open chromatin. This fits with previous observations that variant-free enhancers can reside in cis regulatory domains (CRD) containing functionally interacting enhancers, suggesting that a variant strongly affecting one enhancer within the CRD could have domain-wide effects (Link et al., 2018a). This concept was supported and extended here by HiChIP experiments. In addition to demonstrating that the IL-4 response was primarily associated with pre-existing enhancer-promoter connections, the HiChIP assay also captured a large number of enhancer-enhancer interactions. Examination of these connected enhancers provided evidence that a significant fraction of strain-differential enhancers lacking local variants were connected to strain-differential enhancers containing functional variants. An important future direction will be to further investigate the significance and mechanisms underlying these associations.

Collectively, these studies reveal general mechanisms by which noncoding genetic variation influences signal-dependent enhancer activity, thereby contributing to strain-differential patterns of gene expression and phenotypic diversity. A major future goal will be to incorporate these findings into improved algorithms for prediction of absolute levels and dynamic responses of genes to IL-4 at the level of individual genes.

## Acknowledgments

The authors would like to thank J. Collier and J. Chang for technical assistance, the IGM core for library sequencing, L. Van Ael for assistance with manuscript preparation and Dr. Warren and Dr. Lazarevic for donating *Egr2^flfl^* mice. These studies were supported by NIH grants DK091183 and HL147835 and a Leducq Transatlantic Network grant 16CVD01 to CKG. Sequencing costs were partially supported by DK063491. MAH was supported by a Rubicon grant from the Netherlands Organization for Scientific Research and postdoctoral grants from the Amsterdam Cardiovascular Sciences institute and the American Heart Association.

## Author Contributions

Conceptualization, MAH, ZS and CKG; Formal Analysis, ZS, MAH, IRH and AZ; Investigation, MAH, IC and NJS; Writing, MAH, ZS and CKG; Visualization, ZS, MAH, IRH and IC; Supervision, CKG and MG; Funding Acquisition, CKG.

## Declaration of Interests

The authors declare no conflict of interests.

---

**Figure S1.**
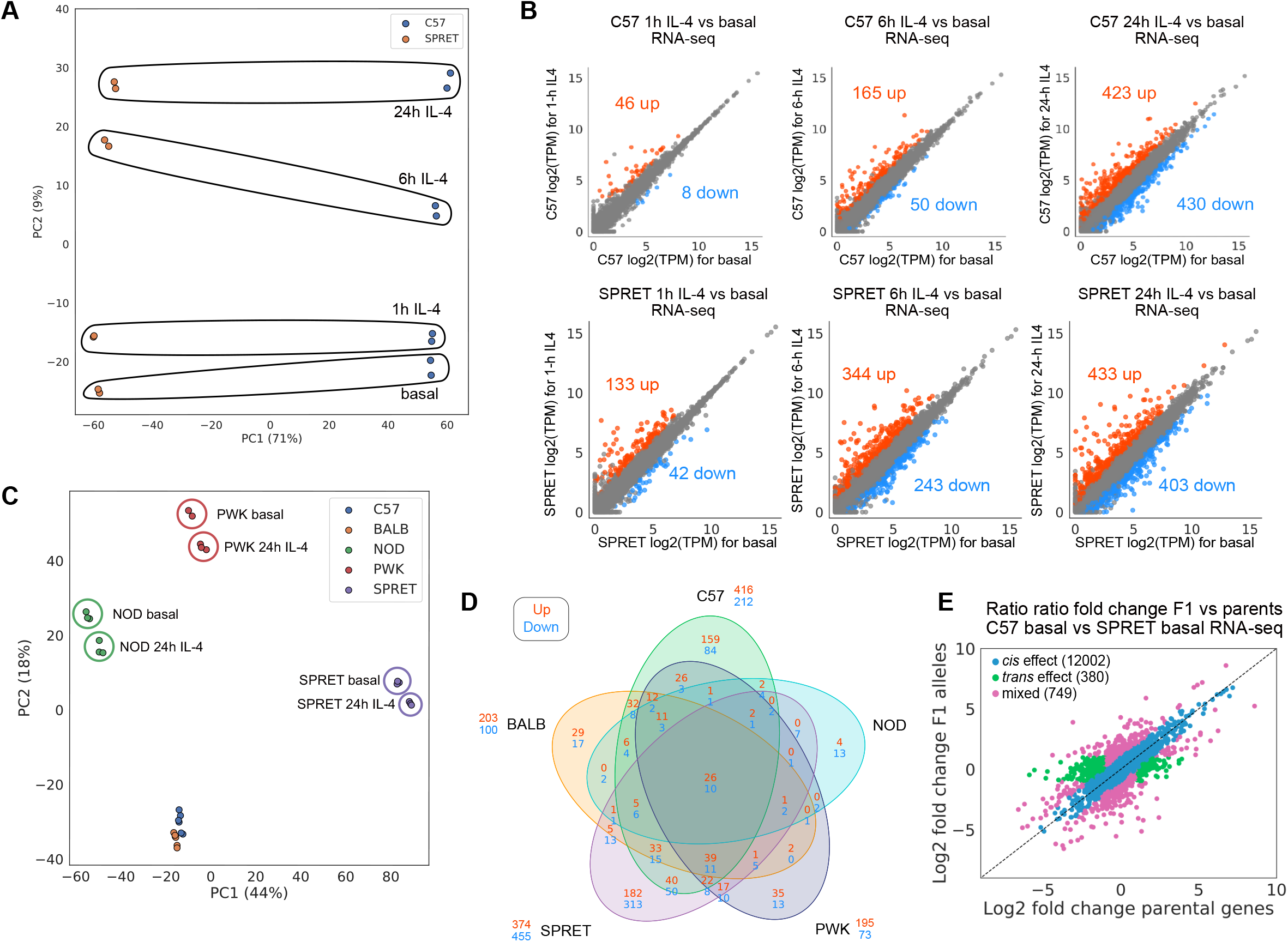
Response to IL-4 is slow and highly divergent in macrophages from different mouse strains. Related to Figure 1. A. PCA plot showing the variance in RNA-seq IL-4 time course data in C57 and SPRET macrophages. B. Scatter plot showing the effects of 1h, 6h and 24h IL-4 stimulation on gene expression in C57 and SPRET BMDMs (n=2 per condition). C. PCA plot showing the variation in macrophages from different strains in response to 24h IL-4. D. Venn diagram of the IL-4 response in macrophages from the five different strains. Repressed and activated genes are plotted that have a twofold change and an q-value<0.05 between untreated and IL-4 stimulated conditions. E. Ratio-ratio fold change plots of allele-specific RNA-seq reads in F1 (C57xSPRET) vs parents C57 or SPRET under basal conditions.

**Figure S2.**
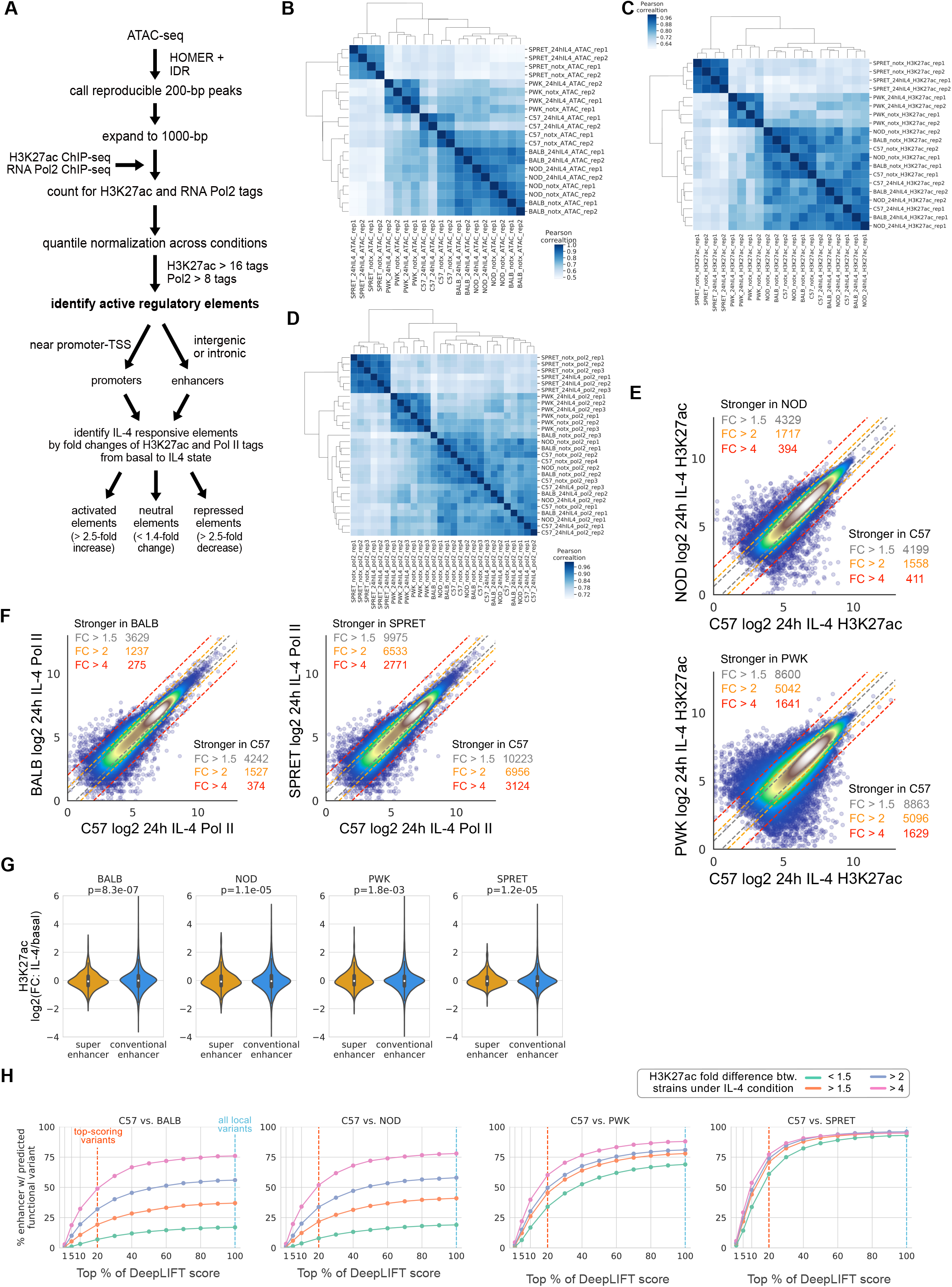
Strain-differential IL-4 induced gene expression is the result of differential IL-4 enhancer activation in macrophages derived from genetically diverse mice. Related to Figure 2. A. Enhancer and promoter selection criteria, including criteria for activated, neutral or repressed elements. B. Clustering of ATAC-seq data in strains macrophages stimulated with IL-4 for 24h. C. Clustering of H3K27Ac ChIP-seq data in strains macrophages stimulated with IL-4 for 24h. D. Clustering of RNApolII ChIP-seq data in strains macrophages stimulated with IL-4 for 24h. E. Comparison of C57 ATAC peaks with H3K27Ac signal to those of NOD or PWK under IL4 treatment conditions F. Comparison of C57 ATAC peaks with RNA Pol2 signal to those of BALB or SPRET under IL4 treatment conditions G. Violin plots showing the difference in H3K27Ac in response to 24h IL-4 between super enhancers and conventional enhancers in BALB, NOD, PWK and SPRET macrophages. Mann–Whitney U test was performed to test the difference between super enhancers and conventional enhancers. H. Percentages of enhancers that contain variants at high-ranked positions based on DeepLIFT scores using different cut-offs. 2G and 2H are based on the top 100% and 20%, respectively.

**Figure S3.**
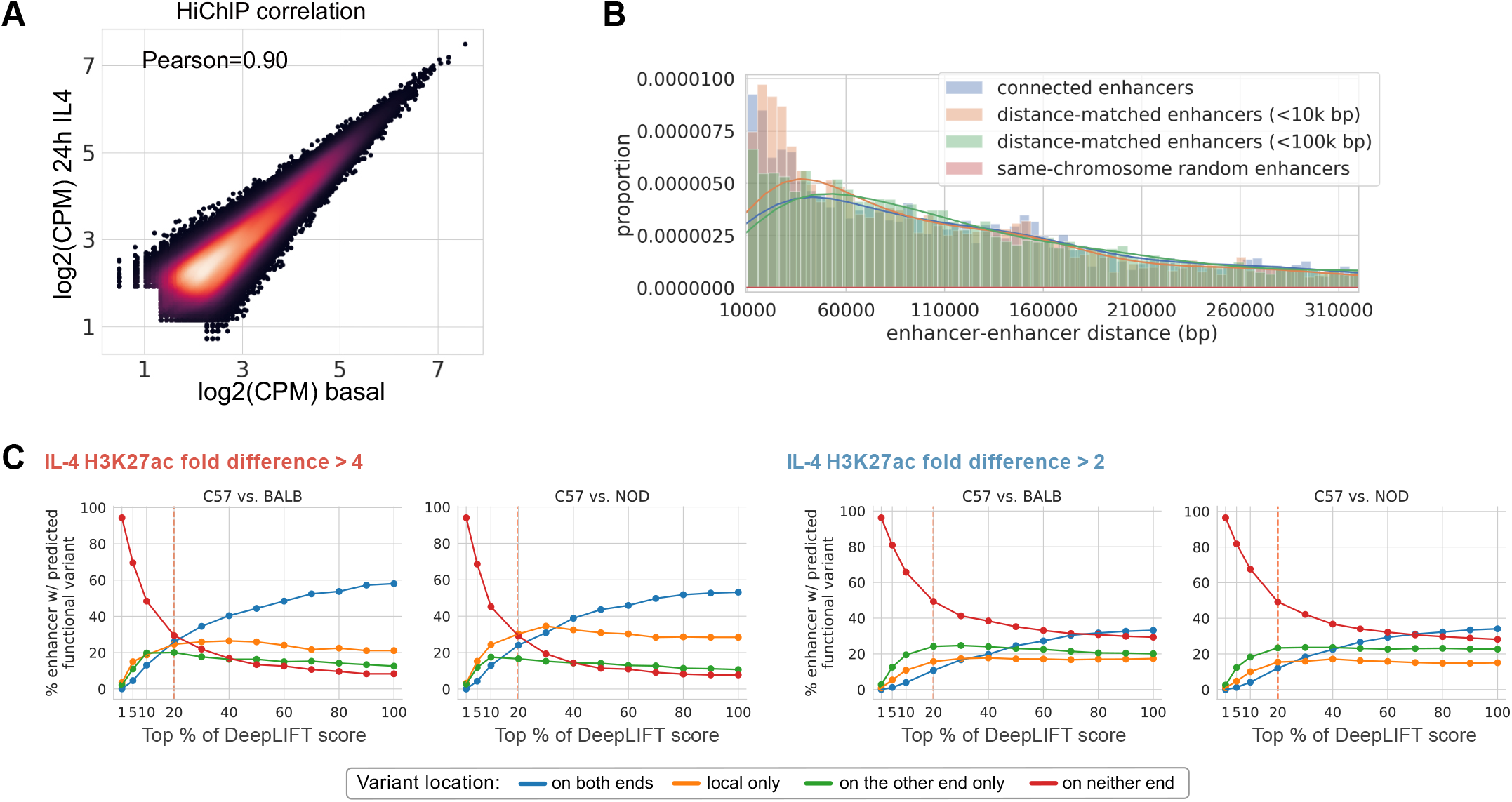
IL-4 enhancers use existent promoter-enhancer interactions to regulate gene activity. Related to Figure 3. A. HiChIP correlation between basal and 24h stimulated C57 macrophages. Each dot represents the amount of reads that connect then bins on both sides of the HiChIP connection in basal and 24h stimulated C57 macrophages. B. Distance distributions of enhancer pairs. Distance-matched random enhancers have similar distances compared to connected enhancers, while the distances between same-chromosome random enhancers are spread out. C. Percentages of interactive enhancers that contain predicted functional variants using different cut-offs for C57 versus BALB and NOD comparisons. 3E is based on the top 20%.

**Figure S4.**
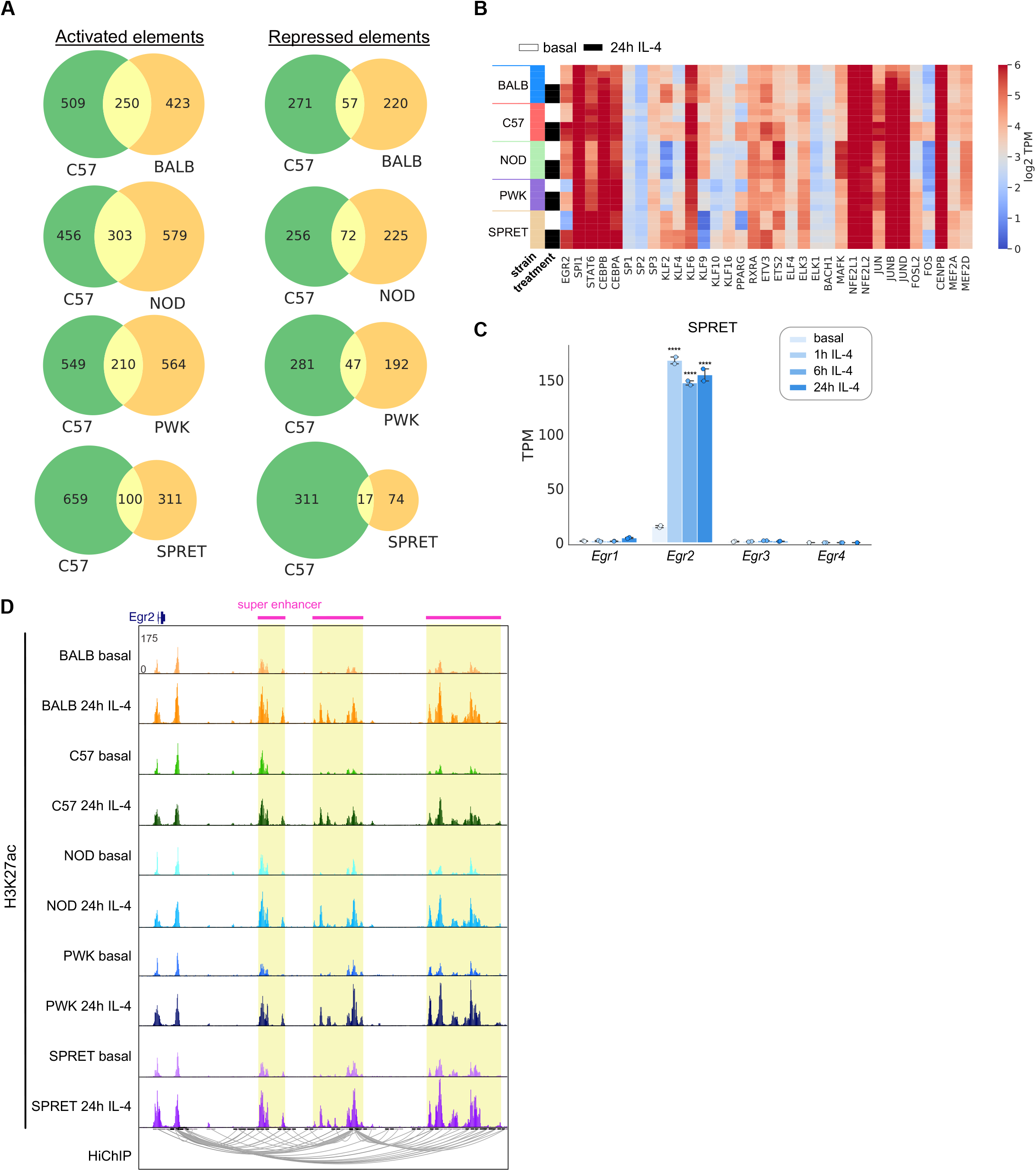
Enhancer mutation motif analysis identifies Egr2 to be strongly associated with late IL-4 enhancer activation. Related to Figure 4. A. Overlap in IL-4 enhancer activation and repression in strain pair-wised comparisons to C57. B. Expression of transcription factors which motifs were in the MAGGIE results (Fig. 4C). C. Gene expression of all Egr family members in SPRET BMDMs under basal conditions and after stimulation with IL-4 for 1h, 6h or 24h. ****q<0.0001, compared to basal. D. Super enhancers of the *Egr2* gene that are strongly conserved in macrophage of the five different strains.

**Figure S5.**
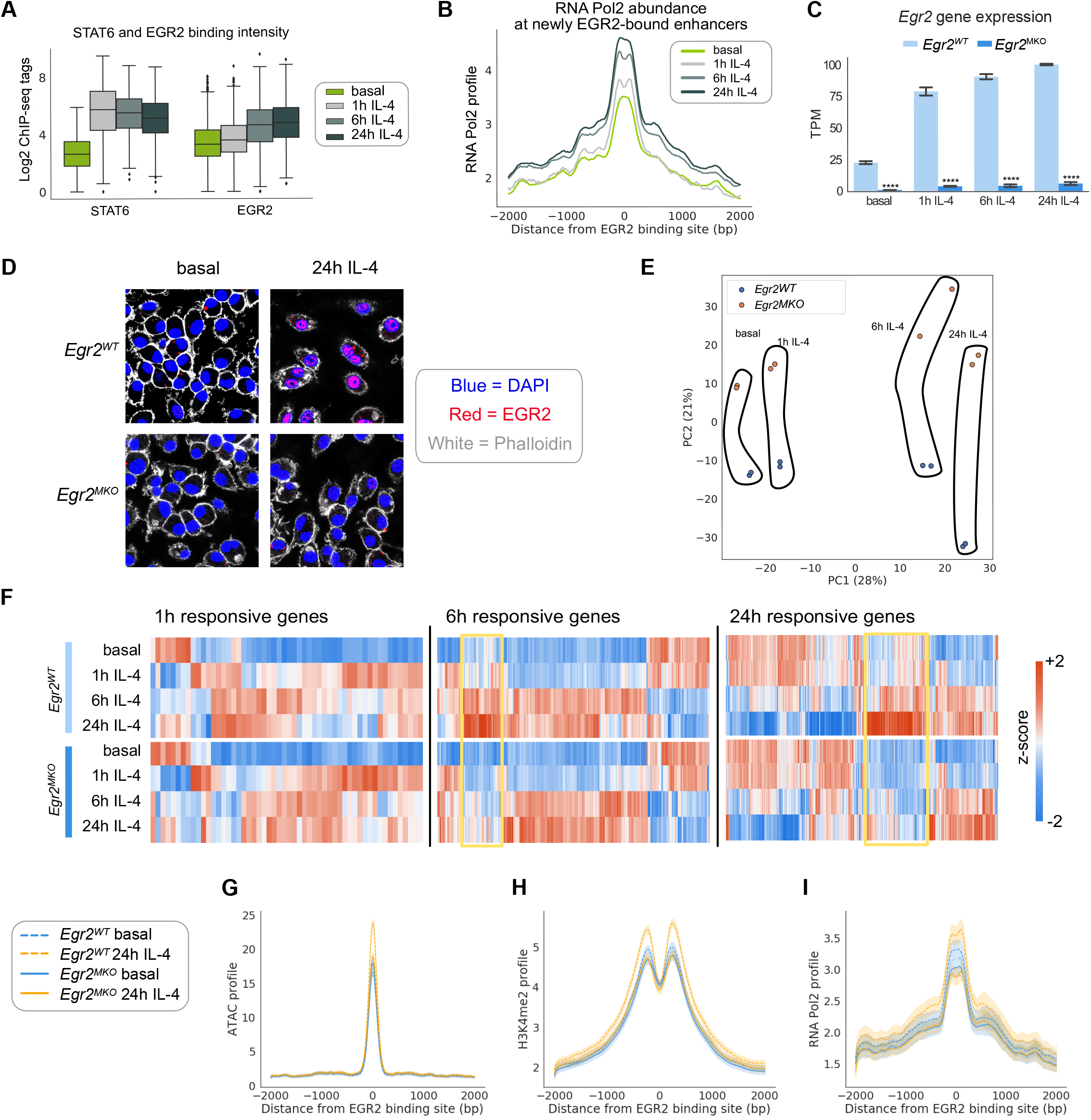
*Egr2* deletion results in decreased IL-4 induced enhancer activation and gene expression. Related to Figure 5. A. STAT6 and EGR2 binding intensity as measured with ChIPseq after IL-4 stimulation over time in C57 BMDMs. B. Enhancer activity as measured with RNA Pol2 binding at 24h IL-4 induced at intergenic and intronic EGR2 peaks in C57 BMDMs. C. Efficient deletion of *Egr2* in *LyzM-Cre*^+^ *Egr2*^fl/fl^ (*Egr2* macrophage knock-out, *Egr2^MKO^)* macrophages, one out of two representative experiments is shown, n=2 per condition. One out of two replicate experiments is shown, ****q<0.0001, compared to *Egr2^WT^* macrophages. D. Immunofluorescence of EGR2 in combination with DAPI and Phalloidin in untreated and IL-4 stimulated *Egr2^WT^* and *Egr2^MKO^* macrophages, one out of two representative experiments is shown. E. PCA plot showing the variance in RNA-seq samples IL-4 time course data in *Egr2^WT^* and *Egr2^MKO^* macrophages. F. Heatmap visualizing the response to IL-4 at different time points in in *Egr2^WT^* and *Egr2^MKO^* macrophages. G. ATAC profile over IL-4 induced EGR2 peaks in *Egr2^WT^*and *Egr2^MKO^* macrophages under basal conditions and after 24h IL-4 stimulation. H. H3K4me2 profile over IL-4 induced EGR2 peaks in *Egr2^WT^*and *Egr2^MKO^* macrophages under basal conditions and after 24h IL-4 stimulation. I. RNA Pol2binding at IL-4 induced EGR2 peaks in *Egr2^WT^*and *Egr2^MKO^* macrophages under basal conditions and after 24h IL-4 stimulation.

**Figure S6.**
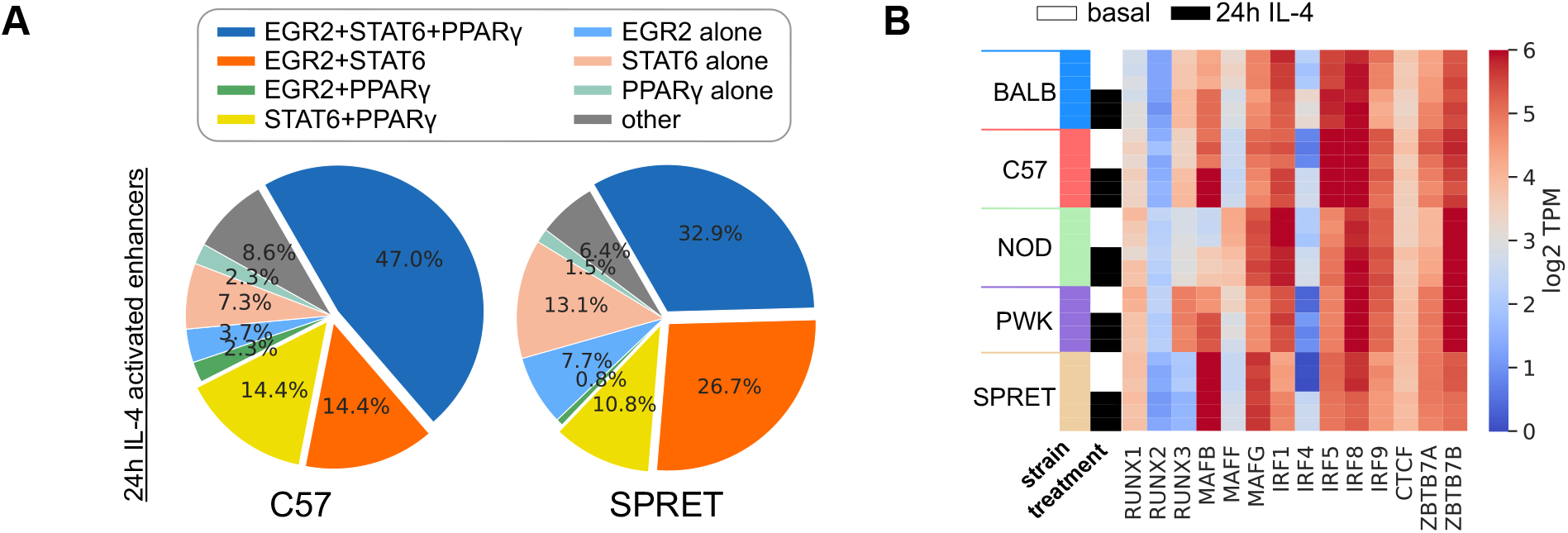
Collaborative and hierarchical transcription factor interactions at IL-4 dependent enhancers. Related to Figure 6. A. Overlap of binding as determined with ChIPseq of the SDTFs STAT6 and PPARg with EGR2 at IL-4 activated enhancers in C57 and SPRET BMDMs. B. Heatmap of gene expression of transcription factors that were in the MAGGIE analysis, Fig. 6D.

**Figure S7.**
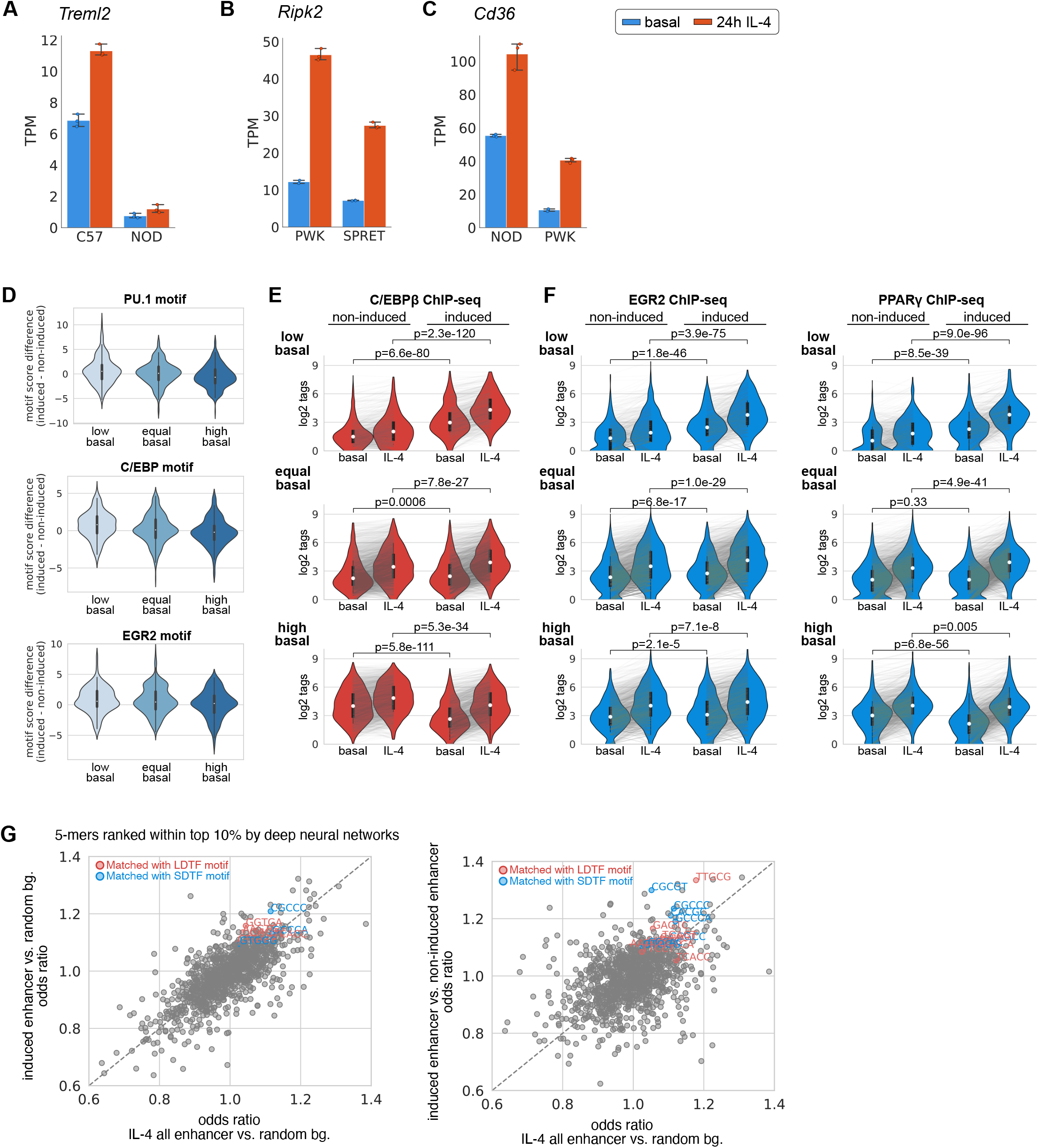
Determinants of absolute levels and dynamic responses of IL-4 responsive enhancers. Related to Figure 7. A *Treml2* gene expression in C57 and NOD macrophages B. *Ripk2* gene expression in PWK and SPRET macrophages C. *Cd36* gene expression in NOD and PWK macrophages D. C/EBPβ binding in non-induced and induced strains in the three different categories of enhancers. E. EGR2 and PPARg binding in non-induced and induced strains in the three different categories of enhancers. F. Score differences of PU.1, C/EBP and EGR2 motifs in the three categories of enhancers. G. Enrichment of 5-mers at top-ranked positions based on different neural network models. 5-mers matched with LDTF (red) or SDTF (blue) motifs based on significant results from TOMTOM are highlighted.

## STAR Methods

### Mice

Female and male breeder mice for C57BL/6J (RRID: IMSR_JAX:000664), BALB/cJ (RRID: MGI:5657790), NOD/ShiLtJ (RRID: IMSR_JAX:001289), PWK/PhJ (RRID: MGI:2678352), and SPRET/EiJ (RRID: MGI:5650926) mice were purchased from Jackson Laboratory. F1 C57 x SPRET (RRID:MGI:5650411) mice were crossed and *Egr2*^fl/fl^ mice were generously donated by dr. Lazarevic and dr. Warren (NIH) and crossed to *LyzM-Cre* mice (Jackson) to achieve myeloid specific targeted deletion of *Egr2*. Mice were housed at the UCSD animal facility on a 12h/12h light/dark cycle with free access to normal chow food and water. All animal procedures were in accordance with University of California San Diego research guidelines for the care and use of laboratory animals. 8-12-week-old healthy female mice were used for all our experiments.

### Bone marrow-derived macrophage (BMDM) culture

Femur, tibia and iliac bones from the different mouse strains were flushed with DMEM high glucose (Corning) and red blood cells were lysed using red blood cell lysis buffer (eBioscience). After counting, 20 million bone marrow cells were seeded per 15cm non-tissue culture plates in DMEM high glucose (50%) with 20% fetal bovine serum (FBS, Omega Biosciences), 30% L929-cell conditioned laboratory-made media (as source of M-CSF, produced as described (Link et al., 2018a) before), 100 U/ml penicillin/streptomycin+L-glutamine (Gibco) and 2.5μg/ml Amphotericin B (HyClone). After 4 days of differentiation, 16.7 ng/ml mouse M-CSF (Shenandoah Biotechnology) was added to the media. After an additional 2 days of culture, adherent cells which were scraped and subsequently seeded onto tissue culture-treated petri dishes in DMEM containing 10% FBS, 100 U/ml penicillin/streptomycin+L-glutamine, 2.5μg/ml Amphotericin B and 16.7 ng/ml M-CSF. Macrophages were left untreated or treated with 20 ng/mL mouse recombinant IL-4 (Peprotech) for 1, 6 or 24 hours.

### Immunofluorescence

Cells were fixed with Cytofix/Cytoperm Buffer (BD, BD554714) for 10 min at room temperature. Cytofix/Cytoperm buffer was removed and cells were washed twice with HBSS containing 2% BSA and 1mm EDTA. Cells were kept in permeabilization/wash buffer (BD, BD554714) for one hour at 4C or until the experiment was performed. Fixed cells were blocked using 3% BSA, 0.1% Triton-PBS for 30 min at room temperature and then with 1/200 of the EGR2 antibody (abcam) overnight at 4C. Next day, cells were washed with 0.1% Triton-PBS, incubated with 1/200 donkey anti-rabbit 555 (ThermoFisher, #A31572) secondary antibody, phalloidin (abcam, ab176759) for staining actin filaments and nuclei were counter-stained with DAPI. After washing with 0.1% Triton-PBS, slides were mounted with Prolong Gold Antifade Reagent (Life Technology, #10144). Images were taken using a Leica SP8 with light deconvolution microscope.

### RNA-seq library preparation

Total RNA was isolated from cells and purified using RNA Directzol micro prep columns and RNase-free DNase digestion according to the manufacturer’s instructions (Zymo Research). Sequencing libraries were prepared in biological replicates from polyA enriched mRNA as previously described (Link, et al. 2018). Libraries were PCR-amplified for 9-14 cycles, size selected using TBE gels or one-sided 0.8X Ampure clean-up, quantified by Qubit dsDNA HS Assay Kit (Thermo Fisher Scientific) and 75bp single-end sequenced on a HiSeq 4000 or NextSeq 500 (Illumina).

### Crosslinking for ChIP-seq

For histone marks, PU.1, C/EBPβ and RNA Pol2 ChIP-seqs, culture media was removed and plates were washed once with PBS and then fixed for 10 minutes with 1% formaldehyde (Thermo Fisher Scientific) in PBS at room temperature and reaction was then quenched by adding glycine (Thermo Fisher Scientific) to 0.125M. For STAT6, PPARg, and EGR2 ChIP-seq, cells were crosslinked for 30 minutes with 2mM DSG (Pierce) in PBS at room temperature. Subsequently cells were fixed for 10 minutes with 1% formaldehyde at room temperature and the reaction was quenched with 0.125M glycine. After fixation, cells were washed once with cold PBS and then scraped into supernatant using a rubber policeman, pelleted for 5 minutes at 400xG at 4°C. Cells were transferred to Eppendorf DNA Lobind tubes and pelleted at 700xG for 5 minutes at 4°C, snap-frozen in liquid nitrogen and stored at −80°C until ready for ChIP-seq protocol preparation.

### Chromatin immunoprecipitation

Chromatin immunoprecipitation (ChIP) was performed in biological replicates as described previously (Seidman et al., 2020). Samples were sonicated using a probe sonicator in 500 μl lysis buffer (10 mM Tris/HCl pH 7.5, 100 mM NaCl, 1 mM EDTA, 0.5mM EGTA, 0.1% deoxycholate, 0.5% sarkozyl, 1 × protease inhibitor cocktail). After sonication, 10% Triton X-100 was added to 1% final concentration and lysates were spun at full speed for 10 minutes. 1% was taken as input DNA, and immunoprecipitation was carried out overnight with 20 μl Protein A Dynabeads (Invitrogen) and 2 μg specific antibodies for PU.1 (Santa Cruz, sc-352X), H3K4me2 (Millipore, 07-030), H3K4me3 (Millipore, 04-745), H3K27ac (Active Motif, 39135), RNA Pol2 (Genetex, GTX102535), STAT6 (Santa Cruz, sc-374021), EGR2 (abcam, ab43020) and C/EBP-β (Santa Cruz, sc-150). Beads were washed three times each with wash buffer I (20mM Tris/HCl, 150mM NaCl, 0.1% SDS, 1% Triton X-100, 2mM EDTA), wash buffer II (10mM Tris/HCl, 250mM LiCl, 1% IGEPAL CA-630, 0.7% Na-deoxycholate, 1mM EDTA), TE 0.2% Triton X-100 and TE 50mM NaCl and subsequently resuspended 25 μl 10 mM Tris/HCl pH 8.0 and 0.05% Tween-20 and sequencing libraries were prepared on the Dynabeads as described below.

For PPAR-g ChIP-seq, fixed cells were lysed in 500 μl RIPA lysis buffer (20 mM Tris/HCl pH7.5, 1 mM EDTA, 0.5 mM EGTA, 0.1% SDS, 0.4% Na-Deoxycholate, 1% NP-40 alternative, 0.5 mM DTT, 1x protease inhibitor cocktail (Sigma)) and chromatin was sheared using a probe sonicator. 1% was taken as input DNA, and immunoprecipitation was carried out overnight with 20 μl Protein A Dynabeads (Invitrogen) and 2 μg of both PPAR-g antibodies (Santa Cruz, sc-271392 and sc-7273). Beads were then collected using a magnet and washed with 175 μl ice cold buffer as indicated by incubating samples on ice for 3 minutes: three times RIPA wash buffer (20 mM Tris/HCl pH7.5, 1 mM EDTA, 0.5 mM EGTA, 0.1% SDS, 0.4% Na-Deoxycholate, 1% NP-40 alternative, 0.5 mM DTT, 1x protease inhibitor cocktail (Sigma)), six times LiCl wash buffer (10 mM Tris/HCl pH7.5, 250mM LiCl, 1 mM EDTA, 0.7% Na-Deoxycholate, 1% NP-40 alternative, 1x protease inhibitor cocktail (Sigma)), twice with TET (10 mM Tris/HCl pH 8.0, 1 mM EDTA, 0.2% Tween-20, 1x protease inhibitor cocktail (Sigma)), and once with TE-NaCl (10 mM Tris/HCl pH 8.0, 0.1 mM EDTA, 50 mM NaCl, 1x protease inhibitor cocktail (Sigma)). Bead complexes were resuspended in 25 μl TT (10 mM Tris/HCl pH 8.0, 0.05% Tween-20) and sequencing libraries were prepared on the Dynabeads as described below.

### ChIP-seq library preparation

ChIP libraries were prepared while bound to Dynabeads using NEBNext Ultra II Library preparation kit (NEB) as previously described (Heinz et al., 2018). DNA was polished, polyA-tailed and ligated after which dual UDI (IDT) or single (Bioo Scientific) barcodes were ligated to it. Libraries were eluted and crosslinks reversed by adding to the 46.5 μl NEB reaction 16 μl water, 4 μl 10% SDS, 4.5 μl 5M NaCl, 3 μl 0.5 M EDTA, 4 μl 0.2M EGTA, 1 μl RNAse (10 mg/ml) and 1 μl 20 mg/ml proteinase K, followed by incubation at 55C for 1 hour and 75C for 30 minutes in a thermal cycler. Dynabeads were removed from the library using a magnet and libraries were cleaned up by adding 2 μl SpeedBeads 3 EDAC (Thermo) in 124 μl 20% PEG 8000/1.5 M NaCl, mixing well, then incubating at room temperature for 10 minutes. SpeedBeads were collected on a magnet and washed two times with 150 μl 80% ethanol for 30 seconds. Beads were collected and ethanol removed following each wash. After the second ethanol wash, beads were air dried and DNA eluted in 12.25 μl 10 mM Tris/HCl pH 8.0 and 0.05% Tween-20. DNA was amplified by PCR for 14 cycles in a 25 μl reaction volume using NEBNext Ultra II PCR master mix and 0.5 μM each Solexa 1GA and Solexa 1GB primers. Libraries were size selected using TBE gels for 200 – 500 bp and DNA eluted using gel diffusion buffer (500 mM ammonium acetate, pH 8.0, 0.1% SDS, 1 mM EDTA, 10 mM magnesium acetate) and purified using ChIP DNA Clean & Concentrator (Zymo Research). Sample concentrations were quantified by Qubit dsDNA HS Assay Kit (Thermo Fisher Scientific) and 75bp single-end sequenced on HiSeq 4000 or NextSeq 500 (Illumina).

### ATAC-seq library preparation

Approximately 80k cells were lysed in 50 μl room temperature ATAC lysis buffer (10 mM Tris-HCl, pH 7.4, 10 mM NaCl, 3 mM MgCl2, 0.1% IGEPAL CA-630), 2.5 μL DNA Tagmentation Enzyme mix (Nextera DNA Library Preparation Kit, Illumina) was added. The mixture was incubated at 37°C for 30 minutes and subsequently purified using the ChIP DNA purification kit (Zymo Research) as described by the manufacturer. DNA was amplified using the Nextera Primer Ad1 and a unique Ad2.n barcoding primers using NEBNext High-Fidelity 2X PCR MM for 8-14 cycles. PCR reactions were size selected using TBE gels for 175 – 350 bp and DNA eluted using gel diffusion buffer (500 mM ammonium acetate, pH 8.0, 0.1% SDS, 1 mM EDTA, 10 mM magnesium acetate) and purified using ChIP DNA Clean & Concentrator (Zymo Research). Samples were quantified by Qubit dsDNA HS Assay Kit (Thermo Fisher Scientific) and 75bp single-end sequenced on HiSeq 4000 or NextSeq 500 (Illumina).

### H3K4me3 HiChIP

For H3K4me3 HiChIP, 10 million formaldehyde crosslinked cells per condition in biological replicates were used. HiChIP was performed as described before (Mumbach et al., 2016). In our experiments, 375 U of MboI (NEB, R0147M) restriction enzyme was used for chromatin digestion. Shearing was performed in three Covaris microtubes per sample and using the following parameters on a Covaris E220 (Fill Level = 6, Duty Cycle = 5, PIP = 140, Cycles/Burst = 200, Time = 200s). H3K4me3 IP was performed using 7.5 μg of antibody (Millipore, 04-745). Final PCR was performed using NEBNext High-Fidelity PCR MM and Nextera general Primer Ad1 and specific Nextera Primer Ad2.n. PCR product was run on a TBE gel (Invitrogen) and libraries were size selected from 250bp to 700bp and cleaned up using 150 ul gel diffusion buffer (500 mM ammonium acetate, pH 8.0, 0.1% SDS, 1 mM EDTA, 10 mM magnesium acetate) and purified using ChIP DNA Clean & Concentrator (Zymo Research). Samples were quantified by Qubit dsDNA HS Assay Kit (Thermo Fisher Scientific) and 75bp paired-end sequenced on a NextSeq 500 (Illumina).

### Data mapping

Custom genomes were generated for BALB/cJ, NOD/ShiLtJ, PWK/PhJ, and SPRET/EiJ mice from the C57BL/6J or mm10 genome as before (Link et al., 2018a) using MMARGE v1.0 (Link et al., 2018b) and the VCF files from the Mouse Genomes Project (Keane et al., 2011). Data generated from different mouse strains were first mapped to their respective genomes using STAR v2.5.3(Dobin et al., 2013) for RNA-seq data, or bowtie2 v2.2.9 (Langmead and Salzberg, 2012) for ATAC-seq, ChIP-seq, and HiChIP data. Then the mapped data was shifted to the mm10 genome using the MMARGE v1.0 ‘shift’ function (Link et al., 2018b) for downstream comparative analyses.

### RNA-seq data analysis

Transcripts were quantified using HOMER v4.11.1 “analyzeRepeats” script (Heinz et al., 2010). TPM values were reported by using the parameters -count exons -condenseGenes -tpm. Log-scaled TPM values were computed by log2(TPM+1). Raw read counts within transcripts were reported by using the parameters -count exons -condenseGenes -noadj. Differentially expressed genes were identified by feeding raw read counts into DESeq2 (Love et al., 2014) through the “getDiffExpression” script of HOMER. IL-4-induced and IL-4-repressed genes were called by fold changes greater than 2 or less than half, respectively, together with p-values smaller than 0.05. Gene ontology analysis was performed using Metascape (Zhou et al., 2019).

Strain-differential genes were defined based on pairwise comparisons between C57 and one of the other strains as being called IL-4-induced or IL-4-repressed in one strain but not in the other. Strain-differential IL-4-induced genes were further classified into three categories based on the relative level of basal expression between the induced strain versus the non-induced strain: high basal, equal basal, and low basal. In the “high basal” group, the non-induced strain has at least 1.5-fold greater basal expression level than the induced strain. The direction of difference flipped for the “low basal” group where the induced strain has over 1.5-fold greater basal expression than the non-induced strain. The genes in between are categorized into the “equal basal” group. RNA-seq data from F1 mice was mapped to both parental genomes (C57 and SPRET) and analyzed in the same way as begore (Link et al., 2018a). In short, the read counts for each transcript were multiplied by the ratio of reads overlapping mutations time 10 and assigned to the parental genomes. Transcripts without any assigned reads in one of the F1 alleles were filtered out. To determine *cis* versus *trans* effects of genetic variation on gene expression, the difference of fold change between parental alleles and F1 alleles were calculated. The genes with majorly *cis* effects were defined by -1 < log2(parental fold change) – log2(F1 fold change) < 1, while those with majorly trans effects were defined by F1 fold change < parental fold change for genes with over +/− 2 fold-change in parental alleles.

### WGCNA analysis

For each strain a differential gene expression analysis was performed to compare IL-4 to basal with Limma Voom (Law et al., 2014). A linear model was fit for all 5 differential comparisons at once, and 1912 genes that were significant with q-value below 0.05 and an absolute fold change of 1.5 in any comparison where included in a Weighted gene co-expression network analysis (WGCNA) (Langfelder and Horvath, 2008). WGCNA was performed with a softpower value of 20, and a signed network was generated. Modules were cut with min module size of 50 and cutheight of 0.999 including PAM-stage. 9 modules were detected of which 2 genes were part of the grey (non-connected) module which was subsequently excluded. Module Eigengenes were calculated and visualized using the verbose-boxplots function that also performed a Kruskall Wallis significance test to test whether all ME values belong to the same distribution and all modules were significantly different between conditions (all P-values below < 0.0012). Two modules exhibited consistent differential expression between IL-4 and notx across strains, while the other 6 modules were most prominently influenced in a strain specific manner. Modules were annotated with Metascape (Zhou et al., 2019).

### ATAC-seq and ChIP-seq data analysis

Based on the HOMER tag directories created from mapped sequencing data, the reproducible ATAC-seq and transcription factor ChIP-seq peaks were identified by using HOMER to call unfiltered 200-bp peaks (parameters -L 0 -C 0 -fdr 0.9 -size 200) and running IDR v2.0.3 on replicates of the same sample with the default parameters (Li et al., 2011). The levels of histone modifications and RNA polymerase II were quantified within +/− 500 bp around the centers of ATAC-seq reproducible peaks using HOMER annotatePeak.pl with parameters “-size -500,500 - norm 1e7”. The transcription factor binding intensities were quantified within +/− 300 bp around the identified ChIP-seq peaks using parameters “-size -150,150 -norm 1e7”. For comparisons across multiple samples (e.g., different time points, mouse strains, transcription factors), we merged the set of peaks first using HOMER mergePeaks “-d given” before quantifying the features above. To visualize the average profile of a dataset around a certain set of peaks, we used HOMER annotatePeaks.pl with parameters “-norm 1e7 -size 4000 -hist 20” to help compute the histograms of 20-bp bins within +/− 2000 bp regions.

### Identification of IL-4 responsive regulatory elements

IL-4 responsive enhancers were identified by the strong fold changes of H3K27ac and RNA Pol2 at intergenic or intronic open chromatin. Reproducible ATAC peaks called from each mouse strain for the basal and IL-4 conditions were first merged and then annotated for genomic positions and the enrichment of H3K27ac and RNA Pol2 within +/− 500 bp using HOMER v4.11.1. Based on the genomic annotations from HOMER annotatePeaks.pl, we classified regions at promoter-TSS as promoters and regions at intergenic or intronic positions as enhancers. Regions with less than 16 normalized tags of H3K27ac or less than 8 normalized tags of RNA Pol2 were filtered out. For the remaining promoters and enhancers, we computed the fold changes of the normalized tags of H3K27ac and RNA Pol2 between basal and IL-4 conditions for each mouse strain. Regions were called IL-4 induced or IL-4 repressed if there were at least 2.5-fold increases or decreases, respectively, from basal to IL-4 state for both histone markers. Regions with less than 1.4-fold changes were called neutral elements.

### Super enhancer

We used ROSE to call super enhancers for the five mouse strains (Whyte et al., 2013). The active enhancers were first merged within each strain for both basal and IL-4 conditions to obtain a set of starting conventional enhancers. Then the ROSE algorithm was run for each strain on the mapped H3K27ac ChIP-seq data with parameter “-t 2500” to exclude TSS. The overall activity of a super enhancer was quantified by the H3K27ac ChIP-seq read counts within the entire identified super enhancer region.

### H3K4me3 HiChIP

#### H3K4me3 ChIP-seq HiChIP reference preprocessing

H3K4me3 ChIP-seqs from basal and 24h IL-4 stimulated macrophages were performed in duplicate with input controls. Fastq files were aligned with bowtie2 (Langmead and Salzberg, 2012) to the mm10 reference genome and peak calling was done with MACS (Zhang et al., 2008) for each replicate separately. Significant peaks were merged using bedtools (Quinlan and Hall, 2010) into a general bed file that was used as corresponding peak-file for MAPS.

#### H3K4me3 HiChIP preprocessing

HiChIP-seq data was processed with MAPS (Juric et al., 2019) at 5000bp resolution as described previously for PLAC-seq (Nott et al., 2019) for all four samples separately, basal and 24h IL-4 duplicate samples combined, and a merge of all four samples.

#### Differential analysis

In order to identify interactions that were significantly stronger in Il4 or control, a differential analysis was performed as described in (Nott et al., 2019). Briefly, significant interactions that were identified in the combined duplicate analysis of IL-4 and notx were merged in a general interaction set. Paired end read counts that fell within these interactions were quantified for each sample separately. The quantified matrix of all significant interactions for all cell types was used as input for Limma (Ritchie et al., 2015) differential interaction analysis. A linear model was fit, with one pairwise contrast (IL4 vs control), with and without batch correction. No interactions were identified that were significantly different between IL4 and control by either method (FDR < 0.1, and absolute log2 FC > 1). Hence, the combined interaction set (generated using both IL4 and control samples) was used for downstream analysis.

### Interactions among promoters and enhancers

Significant interactions captured by HiChIP-seq were overlapped with previously identified active promoters and enhancers for the five mouse strains using HOMER mergePeaks “-d 2500” in order to identify three categories of interactive pairs: enhancer-enhancer, enhancer-promoter, and promoter-promoter. Enhancer-promoter interactions have enhancers on one end and promoters on the other end, while enhancer-enhancer or promoter-promoter interactions are the linked pairs of enhancers or promoters, respectively. We ended up with 242,837 enhancer-enhancer interactions, 247,503 enhancer-promoter interactions, and 73,158 promoter-promoter interactions. To better understand the regulatory landscape associated with IL-4 stimulation, we subsequently focused on enhancer-promoter interactions that contained IL-4 induced, repressed and/or neutral promoters on one end, and IL-4 induced, neutral, and or repressed enhancers on the other end, and quantified the number of interactions between these possible promoter-enhancer combinations in 9 categories as a contingency table. Fisher’s exact test was applied to the contingency table to determine if the any categories were significantly different for three comparisons of interest: IL-4 induced enhancer/promoter interactions vs non-induced enhancer/promoters; IL-4 repressed enhancer/promoter interactions vs non-repressed enhancer/promoters; and IL-4 induced enhancer/promoter interactions vs IL4 repressed enhancer/promoter interactions. For enhancer-enhancer interactions, we pre-selected enhancers that have at least 4-fold difference in H3K27ac ChIP-seq tags between any two strains under the 24h IL-4 condition to obtain a set of strongly strain-differential enhancers. We then computed the Pearson correlation of H3K27ac tags across the five strains for every pair of interactive enhancers among the pre-selected set. The H3K27ac ChIP-seq tags were ordered by C57, BALB, NOD, PWK, and SPRET when calculating correlations. To obtain non-interactive enhancers, we either randomly paired pre-selected enhancers on the same chromosome (same-chromosome) or looked for enhancers within certain distances but not connected based on our data (distance-matched). Distance-matched random enhancers have a distance of their connected enhancers +/− 10 kb or 100 kb.

### Genetic variants at local and connected enhancers

Genetic variation between C57 and the other four strains at strain-differential enhancers was extracted using MMARGE annotate_mutations (Link et al., 2018b), which was based on the VCF files from the Mouse Genomes Project (Keane et al., 2011). Variants were searched within +/150 bp around the centers of enhancers. At least one genetic variant from the comparative strain needs to be present within the search area for such enhancer to be counted as having variants.

### Motif analysis

#### Motif enrichment analysis

Given a certain set of peaks, we used HOMER findMotifsGenome.pl with parameters “-size 200 -mask” to identify de novo motifs and their matched known motifs (Heinz et al., 2010). The background sequences were either the default random sequences or a different set of peaks from a comparative condition in the main text and in the figure legends.

#### Motif mutation analysis

To integrate the genetic variation across mouse strains into motif analysis, we used MAGGIE, which is able to identify functional motifs out of the currently known motifs by testing for the association between motif mutations and the changes in specific epigenomic features (Shen et al., 2020). The known motifs are obtained from the JASPAR database (Fornes et al., 2020). We applied this tool to strain-differential IL-4-responsive enhancers and transcription factor binding sites. Strain-differential IL-4 responsive enhancers were defined as previously described for KLA-responsive enhancers (Shen et al., 2020). In brief, from every pairwise comparison across the five strains, enhancers identified as “IL-4 activated” or “IL-4 repressed” only in one of the compared strains were called strain-differential and were pooled together. For enhancer sites to be included in the analysis, enhancer activity had to be differentially regulated between two strains. As required by MAGGIE, sequences from the genomes of the responsive strains were input as “positive sequences”, and those from the other strains as “negative sequences”. Strain-differential transcription factor binding sites were defined by reproducible ChIP-seq peaks called in one strain but not in the other. “Positive sequences” and “negative sequences” were specified as sequences from the bound and unbound strains, respectively. The output p-values with signs indicating directional associations were averaged for clusters of motifs grouped by a maximum correlation of motif score differences larger than 0.6. Only motif clusters with at least one member showing a corresponding gene expression larger than 2 TPM in BMDMs were shown in figures.

### Categorization of IL-4-induced enhancers

Among the strain-differential IL-4-induced enhancers as described above, we further split them into three categories based on the level of H3K27ac under the basal condition in non-induced strains. “High basal” enhancers have more than 2-fold stronger H3K27ac in non-induced strains, while “low basal” enhancers have more than 2-fold stronger H3K27ac in induced strains (lower basal H3K27ac in non-induced strains). “Equal basal” enhancers are those in between.

### Deep learning

#### Neural network training

We adapted a similar strategy as AgentBind (Zheng et al., 2020) for our training procedure. We implemented a DeepSEA (Zhou and Troyanskaya, 2015) architecture using Keras v2.3.1. DeepSEA consists of three convolutional layers and two fully connected layers. Three models were trained based on our data: IL-4 active enhancers vs. random backgrounds (auROC = 0.894), IL-4 induced enhancers vs. random backgrounds (auROC = 0.919), and IL-4 induced enhancers vs. non-induced enhancers (auROC = 0.796). The enhancer sequences were extended to 300-bp long. In all experiments, we left out sequences on chromosome 8 for cross validation and sequences on chromosome 9 for testing. IL-4 active enhancers and non-induced enhancers were from C57 mice, while IL-4 induced enhancers were pooled from all the five strains in order to reach a comparable sample size. Random genomic backgrounds were generated by randomly selecting nearby GC-matched equal-length sequences on the mm10 genome. We applied binary cross-entropy as the loss function. During each training, the initial learning rate was set as 1e-4 and reduced by a factor of 0.9 when learning stagnated. The training process stopped when the loss value had not decreased for more than 20 epochs.

#### DeepLIFT and importance score

We used DeepLIFT (Shrikumar et al., 2017) to generate importance scores with single-nucleotide resolution using uniform nucleotide backgrounds. For each input sequence, we generated two sets of scores, one for the original sequence and the other for its reverse complement. The final scores were the absolute maximum at each aligned position. We defined predicted functional nucleotides by the top 20% (i.e., top 60) positions within each input 300-bp sequence. To interpret the most important sequence patterns learned by neural networks, we computed the odds ratio of each 5-mer within top 10% of all 5-mers (as in Zheng et al., 2020). Fisher’s Exact test was performed to determine whether 5-mers were enriched. We used TOMTOM (Gupta, et al., 2007) to match 5-mers with known transcription factor binding motifs.

### Data and code availability

All sequencing data have been made available by deposition in the GEO database: GSE159630. The UCSC genome browser (Kent et al., 2002) was used to visualize sequencing data. The codes for neural network model training and interpretation are available on our Github repository: https://github.com/zeyang-shen/macrophage_IL4Response.

## Notes

### Competing Interest Statement

The authors have declared no competing interest.

